# Vectored Immunoprophylaxis and Treatment of SARS-CoV-2 Infection

**DOI:** 10.1101/2023.01.11.523649

**Authors:** Takuya Tada, Belinda M. Dcosta, Julia Minnee, Nathaniel R. Landau

## Abstract

Vectored immunoprophylaxis was first developed as a means to establish engineered immunity to HIV through the use of an adeno-associated viral vector expressing a broadly neutralizing antibody. We have applied this concept to establish long-term prophylaxis against SARS-CoV-2 by adeno-associated and lentiviral vectors expressing a high affinity ACE2 decoy receptor. Administration of decoy-expressing AAV vectors based on AAV2.retro and AAV6.2 by intranasal instillation or intramuscular injection protected mice against high-titered SARS-CoV-2 infection. AAV and lentiviral vectored immunoprophylaxis was durable and active against recent SARS-CoV-2 Omicron subvariants. The AAV vectors were also effective when administered up to 24 hours post-infection. Vectored immunoprophylaxis could be of value for immunocompromised individuals for whom vaccination is not practical and as a means to rapidly establish protection from infection. Unlike monoclonal antibody therapy, the approach is expected to remain active despite continued evolution viral variants.

## Introduction

The concept of vectored immunoprophylaxis was first proposed as an approach to establish protection against HIV infection by the vectored expression of a broadly neutralizing antibody^1^, replacing the need to derive a vaccine immunogen capable of eliciting such antibodies. The approach has since been found to be effective as a therapeutic approach to suppressing virus replication the nonhuman primate SIV model using adeno-associated viruses (AAV) vectors expressing broadly neutralizing antibodies and is currently in clinical trials as a means to suppress HIV replication in infected individuals^2,3^.

Vectored immunoprophylaxis for SARS-CoV-2 through the expression of neutralizing monoclonal antibodies is problematic because of the extraordinarily rapid evolution of the virus. Monoclonal antibody therapy has been highly successful for the treatment of severe COVID-19, decreasing hospitalization and deaths^4^ but has been largely sidelined by the extraordinarily rapid appearance of viral variants that escape neutralization. The first Omicron variant, BA.1, contained 34 mutations in the spike protein, most of which were within or close to the spike protein receptor binding domain and allowed for escape from most of the therapeutic monoclonal antibodies. The Regeneron REGN-COV2 cocktail, a cocktail of REGN10933 and REGN10987 monoclonal antibodies, and the Lilly LY-CoV555 potently neutralize the earlier variants of concern (Alpha, Beta, Gamma and Delta) but their IC_50_s against the Omicron BA.1 variant was greatly increased^5–14^. Vir/GSK VIR-7831 (Sotrovimab) was thought to maintain neutralizing activity against Omicrons BA.1 and BA.2 but was later found to be 10.5- and 340-fold decreased in neutralizing activity against the variants^8–10,12,14,15^. Lilly LY-CoV1404 maintained neutralizing titer against BA.1, BA.2 and BA.4/5^16^ but fails to neutralize the more recent, further mutated Omicron variants BQ1.1 and XBB^17^. The extraordinarily rapid evolution of the virus is likely to continue over the next several years, imposing a challenge to the development of monoclonal antibodies from which the virus cannot escape. The rapidity of virus evolution is also a challenge for the design of effective vaccines.

A strategy to inhibit virus entry that is less subject to escape by novel variants is that of receptor decoys. The strategy is based on soluble forms of the protein, fused to the Fc domain of an immunoglobulin heavy chain to increase its half-life *in vivo^18^*. While viruses can mutate epitopes in the spike protein driven by selective pressure to escape neutralization by antibodies elicited from previous infection or vaccination, the spike protein needs to conserve high affinity binding to its receptor, thereby preserving the neutralizing activity of the receptor decoy. Receptor decoys were first developed as a therapeutic for HIV infection^19,20^. A recombinant protein consisting of the ectodomain of CD4 fused to an immunoglobulin Fc domain was found to bind the viral envelope glycoprotein gp120 with high affinity and potently neutralize the virus *in vitro* but in clinical trials the protein showed no benefit. More recently, the concept was revived by Gardner *et al*. who showed that an enhanced eCD4-Ig protected rhesus macaques from multiple challenges with SIV^3^.

Receptor decoys for SARS-CoV-2 based on soluble forms of ACE2 have been developed by several groups^21–28^. We previously reported the development of a receptor decoy protein termed an “ACE2 microbody” in which the ACE2 ectodomain is fused to the CH3 domain of a human immunoglobulin IgG1 heavy chain Fc region^21^. The decoy proteins, administered by intranasal (i.n.) instillation, have been shown in mouse and hamster models to protect from infection when given shortly prior to infection and to therapeutically suppress virus replication when given up to about 12 hours post-infection^29^. The introduction of point mutations into the ACE2 spike protein binding region of the decoy to increased its affinity for the spike further increased the effectiveness of the proteins^22,24,26^.

Here, we applied vectored immunoprophylaxis to SARS-CoV-2 using AAV and lentiviral vector vectors expressing a modified high affinity ACE2 microbody. AAV2.retro and AAV6.2 vectors, administered either i.n. or by intramuscular (i.m.) injection, provided a high degree of protection in ACE2 transgenic and Balb/c mouse models. The protection was long-lasting and was effective against recent Omicron variants. The AAV vectors were also effective therapeutically when administered shortly post-infection. The lentiviral vector-based decoy was also effective at suppressing virus replication, providing protection that showed no sign of diminishing two months after i.n. administration. Decoy vectored-immunoprophylaxis could be a highly useful means to protect immunocompromised individuals for whom vaccination is less effective and could offer a therapy that remains active against new variants as the emerge.

## Results

### Decoy-expressing AAV vectors inhibit SARS-CoV-2 infection

To determine the feasibility of vectored prophylaxis for SARS-CoV-2, we constructed AAV vectors expressing an ACE2 receptor decoy. The decoy, termed ACE2.1mb, is similar to the ACE2 microbody we previously reported^21^ that consists of the ACE2 ectodomain fused to a single CH3 domain of an IgG1 heavy chain Fc domain **(Figure 1A)**. The protein has been modified by the introduction of point mutations in the ACE2 spike protein binding region that were reported by Chen *et al*. to increase affinity for the spike protein^22^ and by the introduction of an H345A point mutation that inactivates its catalytic activity^30^. The coding sequence was cloned into an AAV vector containing a CAG promoter and virus stock was produced with AAV2.retro and AAV6.2 capsids. AAV2 and AAV6 are reported to have tropism for cells of the mouse and human lung and airway^31–33^. AAV2.retro is a variant of AAV2 that was selected for increased tropism for the central nervous system (CNS) and retrograde movement in axons^34,35^. It has not been reported to transduce lung cells but in pilot experiments, we found that it worked surprisingly well (not shown). AAV6.2 is a variant of AAV6 that contains a single F129L mutation that was found to increase the efficiency of mouse and human airway cell transduction^36^. The ability of the vectors to protect cells from SARS-CoV-2 infection was tested in the lung cell-line A549.ACE2 and the microglial cell-line CHME3.ACE2. The cells were transduced with serial dilutions of the decoy vectors and then challenged 5 days later with D614G, BA.1, BA.2, BA.2.75, BA.4/5 and BQ.1 spike protein-pseudotyped lentiviruses carrying a luciferase reporter genome. At 2 days post-infection (dpi), luciferase activity in the cultures was measured. The results showed that both decoy-expressing AAVs protected A549.ACE2 and CHME3.ACE2 cells from infection (**Figure 1B**). Virus with the D614G spike was the most potently neutralized by the decoy while BA.2 was the most resistant, with a 20-33-fold higher ID_50_ (defined as the multiplicity of infection (MOI) that resulted in a 50% decrease in luciferase activity). The low ID_50_ required to block infection indicates that the decoy was active on bystander cells and that it was not necessary to transduce all of the cells in order to protect the culture.

**Figure 1.**
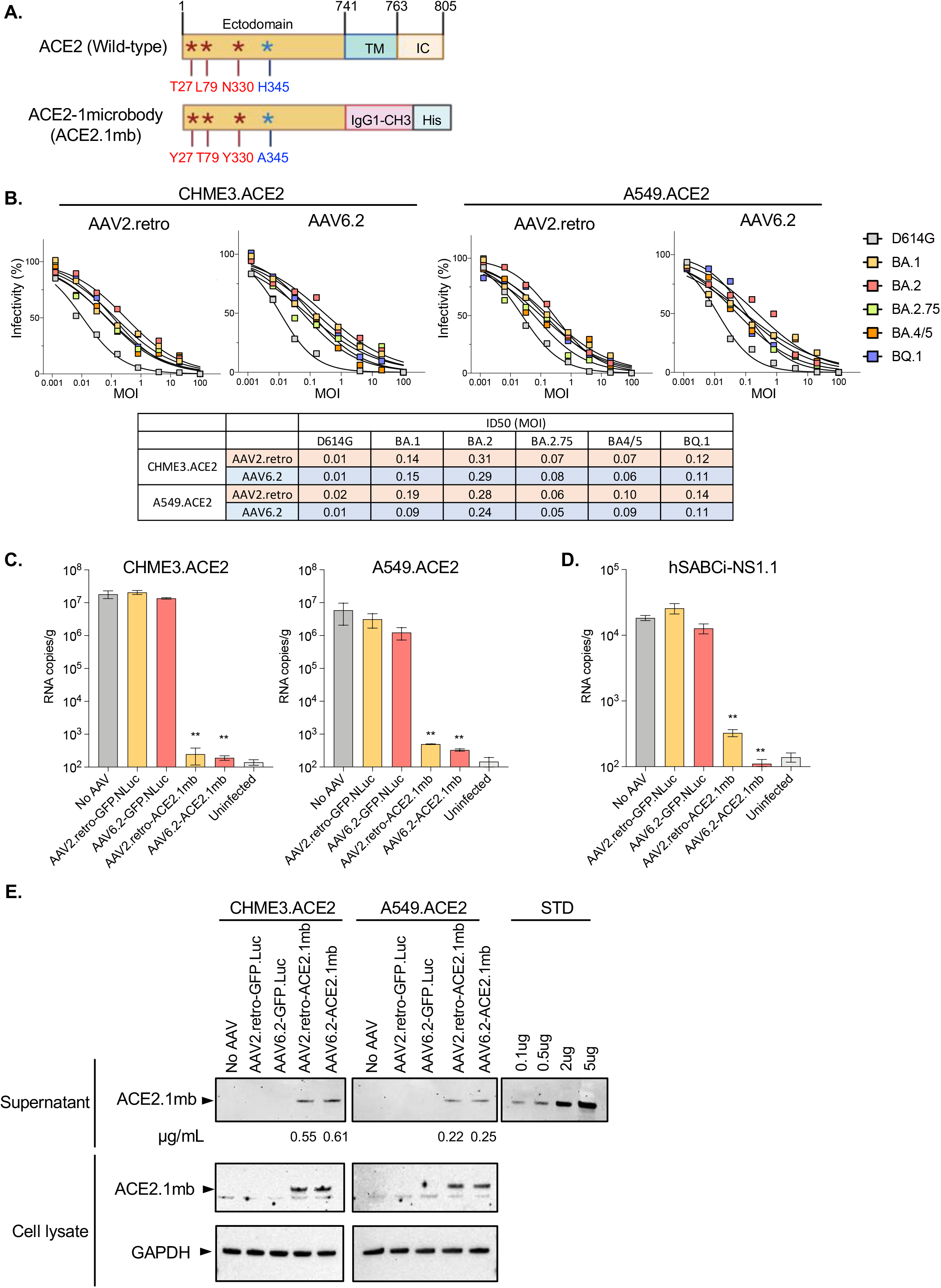
AAV-ACE2.1mb prevents SARS-CoV-2 infection in cell culture. (A) The domain structure of full-length ACE2 is shown above with the ectodomain, transmembrane (TM) and intracellular domain (IC). The structure of the decoy consisting of the ACE2 ectodomain, human IgG1-CH3 and carboxy-terminal His tag. The ACE2 domain contains three high affinity mutations as described by Chan et al. and a H345A mutation in the ACE2 peptidase catalytic site. (B) A549.ACE2 and CHME3.ACE2 cells were transduced with a 5-fold serial dilution of AAV2.retro and AAV6.2 decoy vectors and then challenged with ancestral D614G, Omicron BA.1, BA.2, BA.2.75, BA.4/5 and BQ1 spike protein-pseudotyped lentiviral vectors. Luciferase activity was measured 2-dpi. The curves shown above indicate the infectivity based on luciferase activity normalized to mock vector-transduced cells. Each measurement is shown as the average of duplicates. The table below shows that ID_50_ calculated from the curves shown above. (C) CHME3.ACE2 and A549.ACE2 cells were transduced with AAV2.retro-GFP.nLuc (GFP.Luc), AAV6.2-GFP.nLuc, AAV2.retro-ACE2.1mb, AAV6.2-ACE2.1mb at MOI=0.5. 2 days post transduction, the cells were infected with SARS-CoV-2 WA1/2020 (MOI=0.01). The cultures were lysed 2-dpi and RNA was prepared. Viral RNA copy numbers were determined by RT-qPCR. (D) hSABCi-NS1.1 cells were transduced with AAV2.retro-GFP.nLuc, AAV6.2-GFP.nLuc, AAV2.retro-ACE2.1mb, AAV6.2-ACE2.1mb at MOI=0.5. 2 days post transduction, the cells were infected with SARS-CoV-2 WA1/2020 (MOI=0.01). The cultures were lysed 2-dpi and RNA was prepared. Viral RNA copy numbers were determined by RT-qPCR. Confidence intervals are shown as the mean ± SD. **P ≤ 0.01. The experiment was done twice with similar results. (E) CHME3.ACE2 and A549.ACE2 cells were transduced with AAV2.retro or AAV6.2-ACE2.1mb at MOI=0.5. 3-dpi, secreted decoy protein in the supernatant was pulled-down on NTA beads and bead-bound decoy was detected on an immunoblot probed with His-tag antibody. Pure recombinant decoy protein is shown at right as a standard and was used to determine the amount of protein pulled-down, which is shown below each lane as micrograms decoy pulled-down from 1 ml of culture supernatant. At right is shown decoy protein in the cell lysates is shown below with GAPDH as a loading control.

The ability of the decoy-expressing vectors to inhibit SARS-CoV-2 live virus replication was tested on A549.ACE2, CHME3.ACE2 and hSABCi-NS1.1 cells. The latter is a human small airway basal cell-line grown in air-liquid interface culture conditions and differentiated into mature airway epithelium cell-types the model the respiratory tract. The cells were transduced with decoy-expressing or control GFP.nLuc AAV2.retro and AAV6.2 vectors and challenged a day later with SARS-CoV-2 WA1/2020. Virus replication was measured by RT-qPCR quantification of cell-associated viral RNA copies. The 3 cell-lines supported high levels of SARS-CoV-2 replication (**Figure 1C left)**. Transduction of the CHME3.ACE2 cells with either of the AAV vectors resulted in a 4-5 log decrease in viral RNA, a level that was not significantly higher than uninfected cells. Transduction of the cells by the control AAV had no effect on SARS-CoV-2 replication. The results in the A549.ACE2 cells were similar (**Figure 1C right)**. The vectors were also effective in the hSABCi-NS1.1 human small airway basal cultures although the decrease was less pronounced (50-100-fold) most likely because the cells did not support virus replication as high as in the other cell-lines. The AAV6.2 vector was somewhat more effective than the AAV2.retro vector (**Figure 1D)**. Production of the decoy protein by the transduced CHME3.ACE2 and A549.ACE2 cells was confirmed by pull-down of the protein from the culture supernatant on anti-His tag coated magnetic beads and immunoblot analysis (**Figure 1E)**. The CHME3.ACE2 cells were found to produce about 2-fold more decoy than A549.ACE2 which may have contributed to the greater extent of protection in these cells. The concentration of the decoy protein in the culture medium was 0.2-0.6 μg/ml, a concentration that was greater than the IC_50_ 0.15 μg/ml^21^.

### Vectored immunoprophylaxis *in vivo* by decoy-expressing AAV-vectors

The feasibility of vectored immunoprophylaxis for SARS-CoV-2 with the decoy-expressing AAV vectors was tested in transgenic and non-transgenic mouse models. Decoy-expressing and control GFP AAV2.retro and AAV6.2 vectors were administered to human ACE2 K18 transgenic mice (hACE2 K18 Tg) i.n., i.v. or i.m. After 3 days, the mice were challenged with SARS-CoV-2 WA1/2020 and virus loads in the lung were measured 3-dpi **(Figure 2A)**. The results showed that vector administration i.n. strongly suppressed virus replication in the mice, decreasing the virus load by 5-logs, a level that was indistinguishable from uninfected mice (**Figure 2B**). The control vectors had no effect on virus loads. Histology showed that the lungs of infected untreated mice had prominent signs of interstitial pneumonia with thickened alveolar septa and inflammatory cell infiltration while the lungs of decoy-expressing AAV vectors-treated mice showed no signs of pneumonia and were free of infiltrating inflammatory cells (**Figure 2C**). The lungs of mice treated with the decoy AAV vectors alone in the absence of SARS-CoV-2 infection were clear, indicating that the decoy vectors themselves did not cause pulmonary inflammation (**Figure 2C**). Treatment with the decoy vectors prevented the characteristic loss of body mass associated with untreated SARS-CoV-2 infection (**Figure 2D**). A concern regarding vectored immunoprophylaxis is that the decoy protein or the vectors themselves might induce inflammatory responses in the lungs; however, analysis of proinflammatory and anti-inflammatory cytokine levels (IFNα, IL-10, TNFα, IL12-p70, IL-6 and MCP-1) showed no induction of these cytokines following administration of the vectors (**Figure S1**).

**Figure 2.**
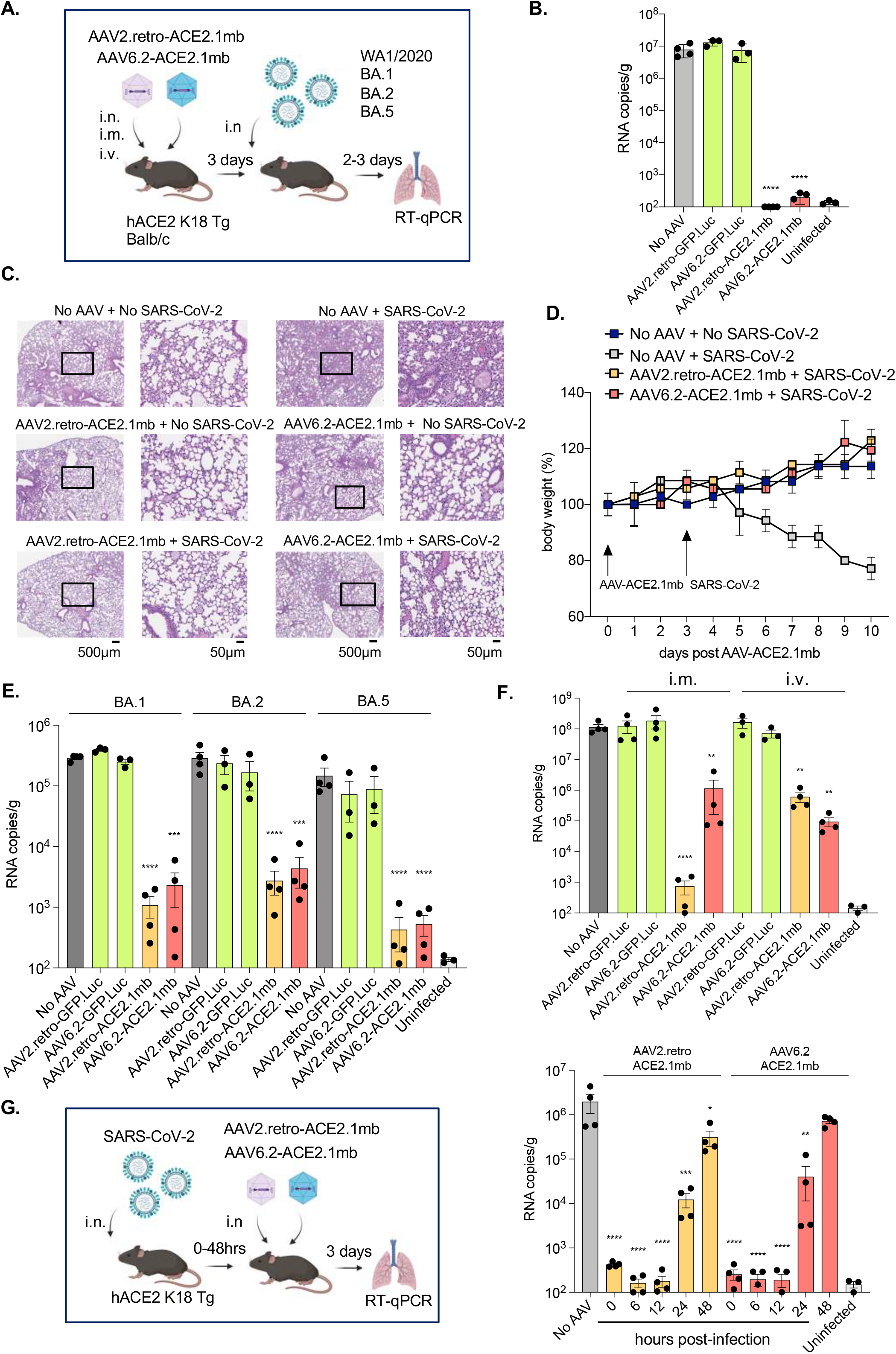
Vectored immunoprophylaxis by decoy-expressing AAV decoy vector and therapeutic use. (A) The experimental scheme for testing decoy prophylaxis is diagrammed. hACE2 K18 Tg and Balb/c mice were treated by i.v. injection or i.n. instillation with decoy-expressing AAV vector or control GFP.nLuc-expressing AAV vector (1 × 10^12^ vg). 3 days post-treatment, the hACE2 K18 Tg mice were challenged with SARS-CoV-2 WA1/2020 and the Balb/c mice were challenged with Omicron BA.1, BA.2 or BA.5 (2 x 10^4^ PFU). Viral RNA copies were quantified 2 days (Omicron) or 3 days (SARS-CoV-2 WA1/2020) post-infection. (B) Mice (n=3-4) were treated with decoy-expressing or control GFP-expressing AAV vectors and challenged with SARS-CoV-2 WA1/2020. Viral RNA copies in the lungs were quantified 3-dpi. (C) H&E stained lung sections from control and decoy-expressing AAV vectors and SARS-CoV-2 infected mice are shown on the left (2 x, scale bars 500μm) and with the boxed area enlarged on the right (20 x, scale bars 50μm). (D) Mice (n=3) were treated with decoy-expressing AAV vectors on day 0 and challenged with SARS-CoV-2 WA1/2020 on day 3. Body weight was measured daily. As controls, the mice were infected with SARS-CoV-2 WA1/2020 but not treated with AAV vector or treated with AAV vector but not infected with SARS-CoV-2 WA1/2020. (E) Balb/c mice (n=3-4) were treated with decoy-expressing or GFP-expressing control AAV vectors and then infected 3 days later with Omicron BA.1, BA.2, or BA.5. Viral RNA copies in the lungs were quantified 2-dpi. (F) Mice (n=4) were administered decoy-expressing AAV vectors i.m. or i.v. and challenged 3-dpi with SARS-CoV-2 WA1/2020. Infected but untreated and uninfected/untreated mice are included as controls. G. Therapeutic use of the decoy-expressing AAV vectors was tested as diagrammed (left). hACE2 K18 Tg (n=4) were infected with SARS-CoV-2 WA1/2020 (2 x 10^4^ PFU) and then treated with decoy-expressing AAVs at time-points up to 48 hours post-infection. Viral RNA in the lung was quantified 3-dpi (right). As controls the mice were untreated (No AAV) or uninfected. Confidence intervals are shown as the mean ± SD. ***P≤0.001, ****P≤0.0001. The experiment was done twice with similar results.

To test the effectiveness of the decoy vectors in protecting against the Omicron variants, Balb/c mice, which support high level replication of SARS-CoV-2 Omicron variants through the endogenous murine ACE2^37,38^, were treated i.n. with decoy-expressing AAV or control AAV2.retro or AAV6.2 vector and then challenged with Omicrons BA.1, BA.2 or BA.5. The results showed that i.n. administration of decoy-expressing AAV vectors caused a dramatic decrease in virus loads as compared to the control vectors (**Figure 2E**). The decoy-expressing vectors were most effective against the BA.5 variant, decreasing the virus load 1,000-fold and least effective against BA.2, decreasing virus load 100-fold (**Figure 2E**), a pattern that was similar to what was found with the pseudotyped lentiviruses *in vitro*. Both AAV vectors were effective although the AAV2.retro seemed to be slightly more suppressive against all three Omicrons. This conclusion was confirmed in a dose-response analysis which showed that the decoy-expressing AAV2.retro vector was about 10-fold more effective at virus load suppression. A 10,000-fold decrease in virus load required 1 × 10^10^ vector genomes (vg) for AAV2.retro. The same degree of suppression by AAV6.2 required 1 × 10^11^ vg (**Figure S2**).

Administration of the vectors i.n. delivered the vectors to the relevant organ but it was possible that delivery by routes that targeted a different site in the body might also be effective given that the decoy protein is stable *in vivo* and freely diffusible^29^. In support of this approach, i.m. delivery of AAV-vectored immunoprophylaxis was effective for the suppression of SIV replication in the macaque model^39^. We therefore tested the effectiveness of i.v. and i.m. administration of the decoy-expressing AAV vectors **(Figure 2A)**. The results showed that i.m. administration was highly effective, decreasing the virus load by 5-logs compared to control vector, a level that was indistinguishable from uninfected mice (**Figure 2F**). I.v. administration was much less effective, decreasing the virus load by only a 2-logs.

### Therapeutic use of vectored immunoprophylaxis for SARS-CoV-2

The studies described above tested the prophylactic effect of the decoy-expressing AAVs administered prior to SARS-CoV-2 infection. It was possible that the approach might also be effective therapeutically by administration post-infection. The effectiveness of administration post-infection would depend on how soon post-infection they were administered and how fast the vectors transduced lung cells and produced the encoded protein to establish an inhibitory concentration in the respiratory tract. To determine this, we infected mice with SARS-CoV-2 and then treated them at increasing times post-infection **(Figure 2G)**. The results showed that the decoy-expressing vectors were effective when administered concomitant with SARS-CoV-2 and up to 12 hours post-infection (**Figure 2G**). The treatment was partially effective at 24-hours and lost efficacy at 48-hours. The results demonstrate remarkably rapid transduction and biosynthesis of the decoy protein by the AAV vectors. While this time course would appear to be too short to be of therapeutic use, the kinetics closely mirror what is seen in monoclonal antibody therapy of SARS-CoV-2 in mouse models^1111118^ suggesting that in humans, where the time-course of disease is slower, the AAV vectors might act with the kinetics similar to that of highly effective monoclonal antibodies.

### AAV decoy-expressing vectored immunoprophylaxis is highly durable

Although AAV does not integrate at a significant frequency into the host cell genome, the genome remains stable in the host cell. In nonhuman primates and in clinical trials, AAV vectors have been shown to maintain long-term expression of an encoded gene *in vivo^40^*. To test the durability of the decoy-expressing AAV vectors, we constructed AAV2.retro and AAV6.2 vectors that expressed a decoy-luciferase fusion protein. The vectors were administered i.n. to mice and the mice were live-imaged over the next 30 days. Expression by both vectors in the lungs was first detected 24 hours post-treatment and then increased to maximal by day 3 **(Figure 3A)**. Expression levels remained stable through day 14 after which they decreased slightly by day 30. Measurement of luciferase activity in tissue homogenates further demonstrated durable expression by the vectors (**Figure 3B**). The decoy proteins were readily detectable on day 1 and the following day, expression increased 25-fold. To determine the durability of viral load suppression by the vectors, mice were treated and then challenged with SARS-CoV-2 over a 30-day period. The results showed that the decoy-expressing vectors strongly suppressed the virus loads of mice infected that had been infected up to 30-days post-treatment (**Figure 3C**). Virus load suppression appeared to begin to wane 30 days post-treatment but was still highly active, with the AAV2.retro vector suppressing virus load nearly 1000-fold. The results were consistent with the slight decrease in *in vivo* expression levels found for the decoy-luciferase fusion protein.

**Figure 3.**
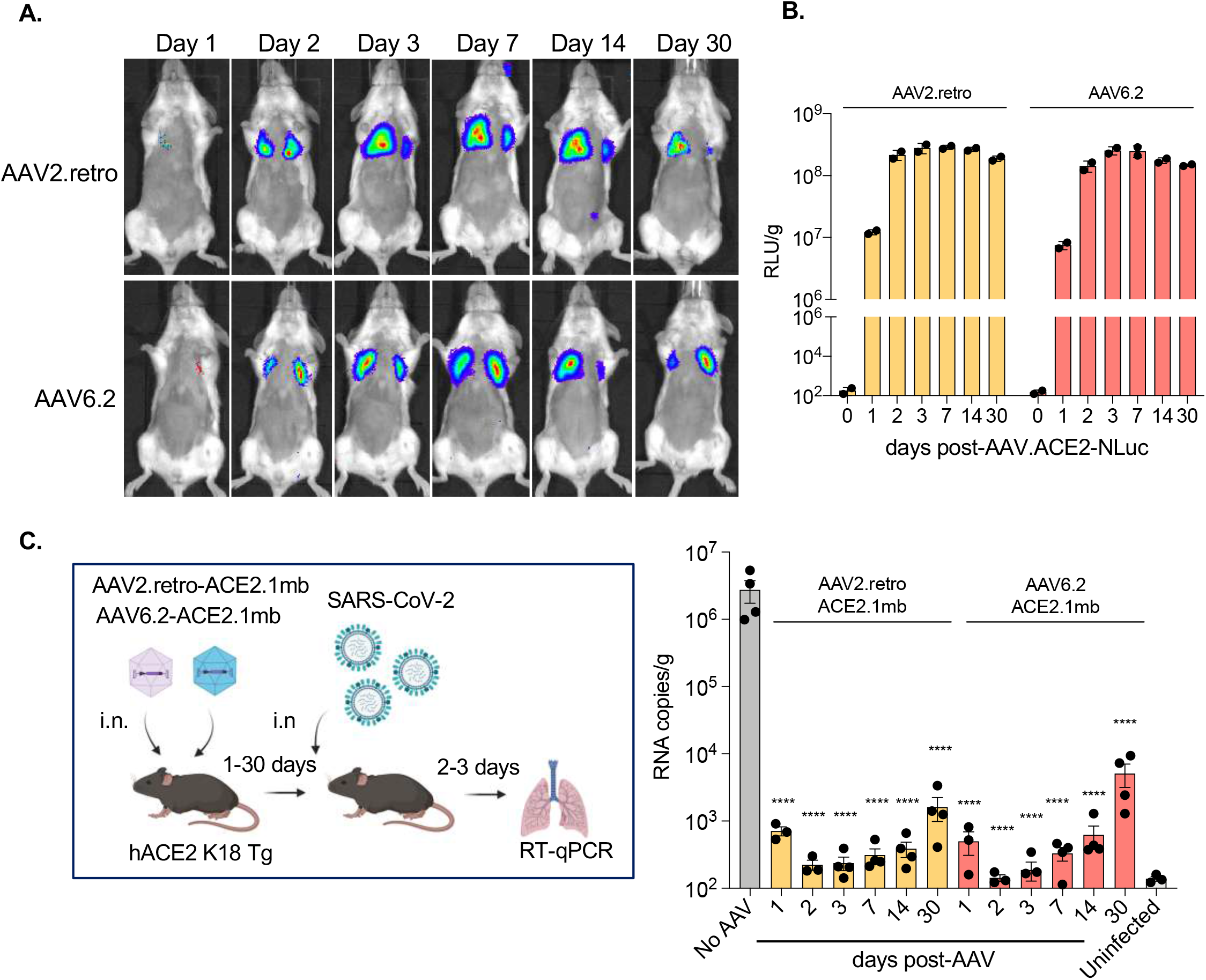
Durable vectored immunoprophylaxis by decoy-expressing AAV vectors. (A) Balb/c mice (n=3) were injected i.n. with decoy-luciferase fusion protein-expressing AAV vectors (1 × 10^12^ vg). Luciferase activity was visualized by live imaging over 30 days at the indicated time points. Representative images of a mouse from each group are shown. (B) Luciferase activity in lung tissue homogenates from mice treated with the decoy-luciferase expressing AAV vectors (n=2) was measured over the time course. (C) The experimental scheme to test the durability of AAV vectored immunoprophylaxis is diagrammed (left). hACE2 K18 Tg (n=4) were injected with AAV decoy. At 1-, 2-, 3-dpi, the mice were challenged with SARS-CoV-2 (2 x 10^4^ PFU) and viral RNA in the lungs was quantified. The results are shown as a histogram (right). SARS-CoV-2 infected/AAV untreated (No AAV) and AAV untreated/SARS-CoV-2 uninfected (Uninfected) controls are shown. Confidence intervals are shown as the mean ± SD. ****P≤0.0001. The experiment was done twice with similar results.

### Increased durability of vectored immunoprophylaxis with a decoy-expressing lentiviral vector

The waning of protection established by the AAV vectors at 30 days led us to test whether a different vector might be able to extend the durability protection. The use of an alternative vector was also of interest in light of concerns about the possibly of pre-existing immunity to AAV in some individuals noted in clinical trials^41^. Lentiviral vectors are generally not subject to pre-existing immunity in humans. Moreover, because lentiviruses integrate into the host cell genome, the vectors are maintained stably in the cell and in daughter cells that may be generated, allowing for the possibility of long-term decoy expression and increased durability of protection. In addition, pseudotyping of the vectors by VSV-G results in a broad target cell tropism. To test the feasibility of lentiviral vectored immunoprophylaxis, we constructed a decoy-expressing lentiviral vector and compared its effectiveness to the AAV vectors. Transduction of A549.ACE2 and CHME3.ACE2 cells with the vector showed that it expressed the decoy protein at a level similar to those of the AAV vectors as measured in the supernatant pull-down assay (**Figure S3**). To determine the potency of the protection, the cell-lines were transduced with a serial of the decoy-expressing lentiviral vector and then challenged with the D614G and Omicron spike protein-pseudotyped reporter viruses. The results showed that vector was highly protective against all of the variants (**Figure 4A**). Overall, the potency of virus neutralization, calculated by the MOI required to decrease infection by 50%, was very similar to that of the AAV vectors. As for the AAV vectors, BA.2 was the most resistant to neutralization (8-fold in CHME3.ACE2 and 5.8-fold in A549.ACE2) (**Figure 4A, below**). The ability of the vectors to neutralize the viruses at low MOIs confirmed that only a small fraction of the cells needed to be transduced to protect the entire population.

**Figure 4.**
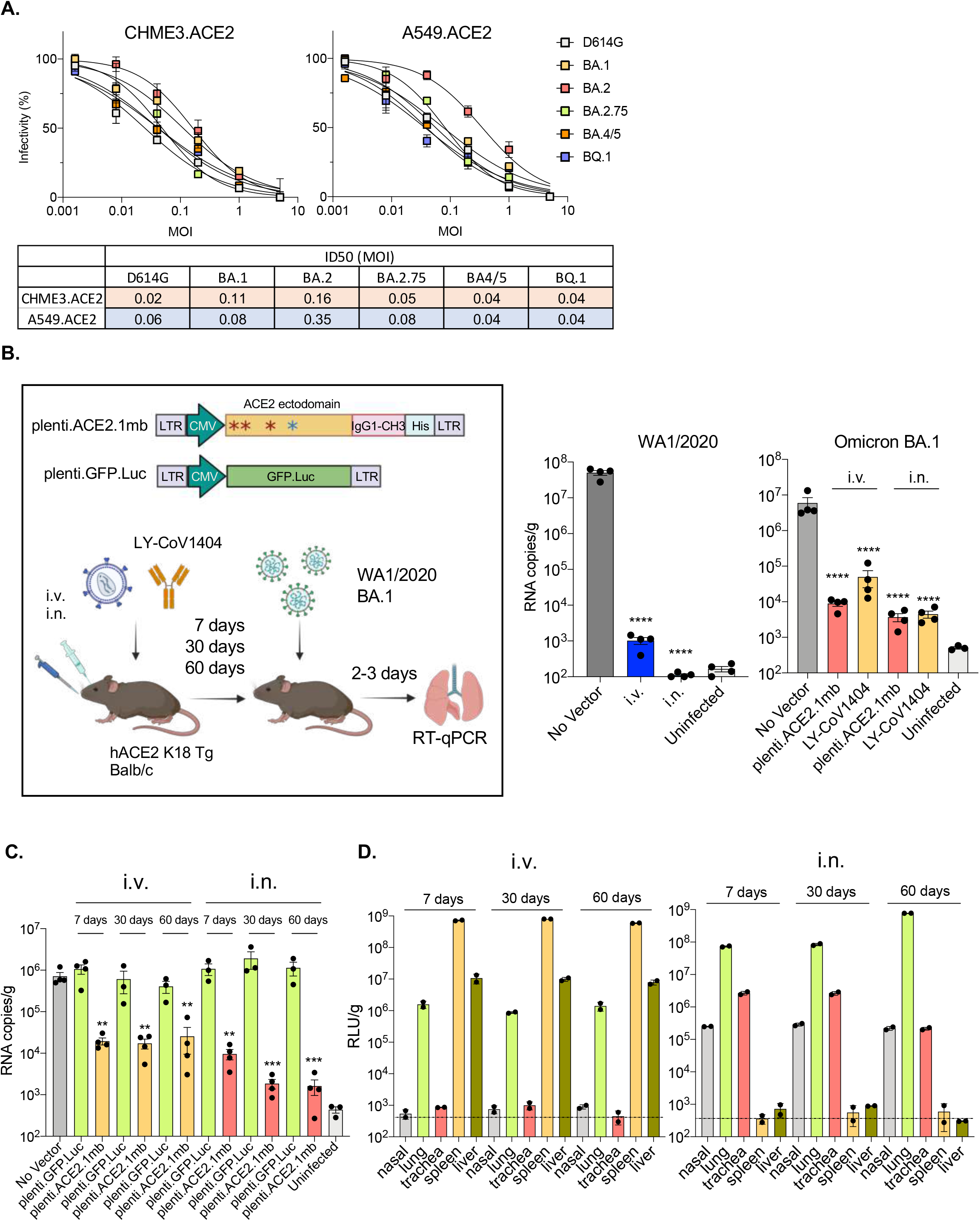
Long-term vectored immunoprophylaxis by decoy-expressing lentiviral vector. (A) A549.ACE2 and CHME3.ACE2 cells were transduced with a 5-fold serial dilution of decoy-expressing lentiviral vectors and then challenged with D614G, Omicron BA.1, BA.2, 2.75, BA.4/5 and BQ.1 spike protein-pseudotyped lentiviral vectors. Luciferase activity was measured 2-dpi (above). The curves indicate infectivity based on luciferase activity normalized to mock vector-transduced cells. Measurements are the average of duplicates. ID_50_s calculated from the curves are shown in the table (below). (B) Structure of lentiviral vector and experimental scheme are shown. hACE2 K18 Tg mice or Balb/c were injected with lentiviral vector (5 x 10^6^ IU) i.p., i.v. or i.n. injection. One week later, the mice were challenged with 2 x 10^4^ PFU of SARS-CoV-2 WA1/2020 (hACE2 K18 Tg) or Omicron (Omicron). Viral RNA in the lungs was quantified 3-dpi. (C) Mice were administered luciferase-expressing lentiviral vector i.v. or i.n. (n=2). Tissues (nasal, lung, trachea, spleen and liver) were harvested and luciferase activity was measured over 60 days at the indicated time-points. (D) Decoy-expressing lentiviral vectors (5 x 10^6^ IU) were administered i.v. or i.n. and after 7-, 30- and 60-days challenged with SARS-CoV-2 (2 x 10^4^ PFU). Viral RNA was quantified 3-dpi. Confidence intervals are shown as the mean ± SD. **P ≤ 0.01, ****P≤0.0001. The experiment was done twice with similar results.

To determine whether the decoy-expressing lentiviral vector could establish vectored immunoprophylaxis, the vector was administered i.v. or i.n. and after 7 days, the mice were challenged with WA1/2020 or Omicron BA.1. Virus loads in the lung were quantified 3-dpi (**Figure 4B, left**). In mice challenged with WA1/2020, i.v. injection resulted in a nearly 5-log decrease in virus load while administration i.n. further decreased the virus load to undetectable levels **(Figure 4B, middle)**. The analysis of proinflammatory and anti-inflammatory cytokine levels (IFNa, IL-10, TNFa, IL12-p70, IL-6 and MCP-1) showed that administration of the lentiviral vector had no significant effect on the levels of these cytokines (**Figure S1**). A dose-response analysis in which mice were administered decreasing amounts of the vector i.v. or i.n. confirmed the great efficacy of i.n. administration; at a dose of 1 × 10^6^ IU, virus loads were 100-fold lower in mice treated i.n. compared to i.v. (**Figure S4**). Treatment with the vector also protected against Omicron BA.1 but did not suppress virus replication to as great an extent, resulting in low level virus replication in mice treated i.v. or i.n. To compare the protective effect of the vector with that of a therapeutic monoclonal antibody, mice were administered the highly potent neutralizing monoclonal antibody LY-CoV1404 by i.v. and i.n. routes and then infected with the Omicron BA.1 variant. The results showed that the monoclonal antibody decreased the virus loads more effectively i.n. than i.v. but was not as effective as the lentiviral vector **(Figure 4B, right)**. To test the durability of lentiviral vectored immunoprophylaxis, mice were treated i.v. or i.n. with decoy-expressing or control GFP-expressing lentiviral vector and challenged 7, 30 and 60 days later with SARS-CoV-2 WA1/2020. The protection was found to persist over the 60-day time-course (**Figure 4C**). I.n. administration of the vector caused a greater decrease in virus loads which interestingly, became even more pronounced over time.

To understand the basis of the long-lasting protection provided by lentiviral vectored immunoprophylaxis, we administered a GFP/luciferase-expressing lentiviral vector i.v. or i.n. and determined the level of expression over the 60-day time-course by measuring luciferase activity in cell lysates prepared from different tissues. The results showed that i.v administration resulted in high level expression in the spleen at day-7 and moderate expression in the lungs and liver (400-fold less in lung and 50-fold less in liver). The expression levels remained constant over the time-course **(Figure 4D)**. There was no detectable expression in nasal tissue and trachea. Administration of the vector i.n. resulted in high level expression in the lung, moderate levels in the trachea (about 30-fold less on day 7) and nasal tissue. Levels in the spleen and liver were undetectable. Expression levels remained constant at 30-days. At 60-days, expression in the lung increased about 8-fold, a finding that could explain the increase in virus load suppression at this time-point in mice treated by i.n. administration of the vector **(Figure 4C)**.

### Comparison of lung cell-types transduced by the AAV and lentiviral vectors

The effectiveness and longevity of the vectored immunoprophylaxis depends both upon the cell-types and half-lives of the cells transduced by the vectors. To understand the basis of durable protection, we characterized the cell-types transduced in the lung by the AAV and lentiviral vectors. Mice were administered GFP-expressing AAV and lentiviral vectors i.n. and the GFP+ cells. The lungs were harvested 3 days later and the cells disaggregated. The cells were then analyzed by flow cytometry using antibodies that distinguished various pulmonary cell-types. The results showed that the majority of the cells transduced in the lungs by AAV2.retro and AAV6.2 were epithelial (79.5% and 94.2%, respectively) **(Figure 5A)**. Of the cells transduced by AAV2.retro, 20.5% were leukocytes while AAV6.2 transduced fewer leukocytes (5.8%). Analysis of the transduced leukocytes showed that the majority of cells were interstitial macrophages and neutrophils with smaller proportions of T cells, B cells, DCs, monocytes and alveolar macrophages. The distribution of leukocytes transduced by AAV6.2 was roughly similar. The lentiviral vector targeted a larger proportion of leukocytes (57%). Of the transduced leukocytes, the greatest proportion were DCs (26.3%) with substantial contributions from B cells (20.7%) and monocytes (18.5%) **(Figure 5B)**. It is possible that the long-lasting expression by the lentiviral vector resulted from the increased transduction of leukocytes, particularly of the DCs, as these cells are thought to be long-lived residents in the lung^42^.

**Figure 5.**
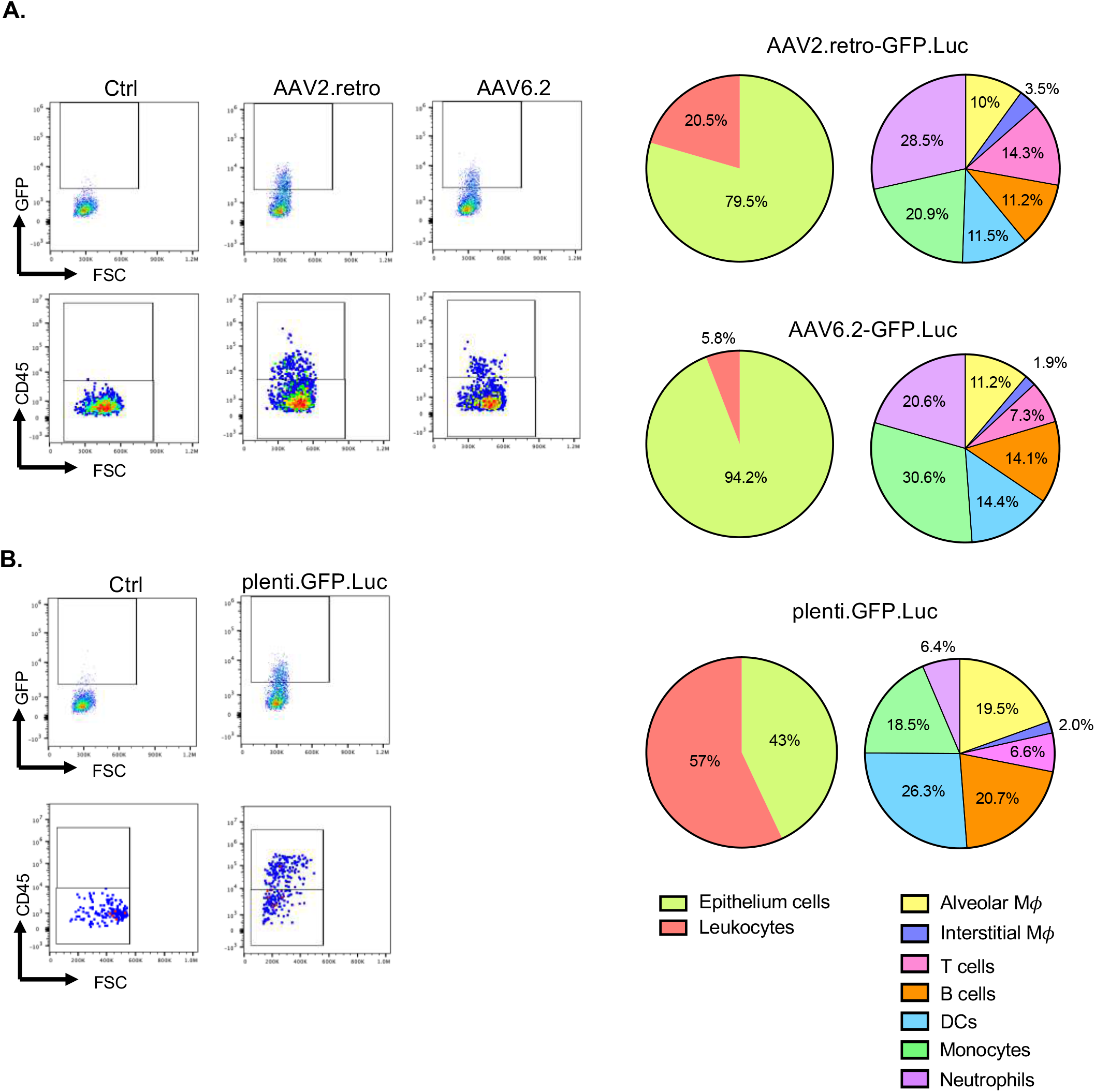
Comparison of lung cell subpopulations transduced by AAV and lentiviral vectors. (A) GFP-expressing AAV vectors were administrated i.n. After 3 days, the lungs were harvested and the tissue were enzymatically disaggregated. The cells were analyzed by multi-color flow cytometry with cell-type specific marker antibodies to distinguish subpopulations defined as follows: Leukocytes (CD45+), epithelial (CD45-), alveolar macrophages (CD45+, F4/80+, SiglecF+), interstitial macrophages (CD45+, F4/80+, SiglecF-), DCs (CD45+, F4/80-, CD11c+), T cells (CD45+, CD3+), B cells (CD45+, CD19+), monocytes (CD45+, CD11b+, CD14+), neutrophils (CD45+, CD62L+, Ly6C/Ly6G+). Representative flow cytometry plots of the GFP+ cells and GFP+/CD45+ populations are shown on the left and the subpopulations within the GFP+/CD45+ leukocytes are shown in the pie charts on the right. (B) Mice were administered GFP-expressing lentiviral vector i.n. GFP+ cells in the lung were analyzed by flow cytometry as in (A). Representative flow cytometry plots of the GFP+ cells and GFP+/CD45+ populations are shown on the left and the subpopulations within the GFP+/CD45+ leukocytes are shown in the pie charts on the right.

## Discussion

Vectored prophylaxis was first developed as an approach to protect against HIV infection^1^ in which broadly neutralizing antibody was expressed in an AAV vector and was later expanded to the use of an enhanced CD4-Ig fusion protein that established a high degree of resistance to SIV infection in treated macaques^2,3^. We report here that that vectored expression of a high affinity ACE2 microbody protein in which the ectodomain of ACE2 was mutated to increase its affinity for the viral spike protein and inactivate catalytic activity fused to the CH3 domain of an IgG heavy chain Fc^21^ established a high degree of protection from SARS-CoV-2 in mouse models. Decoy-expressing AAV2.retro and AAV6.2 vectors were both highly effective at establishing vectored immunoprophylaxis in ACE2 K18 Tg and Balb/c mice. Mice treated with the decoy-expressing AAV2.retro and AAV6.2 vectors were highly resistant to SARS-CoV-2 infection. Upon challenge with high titered SARS-CoV-2 WA1/2020, viral RNA in the lungs 3-dpi was undetectable, corresponding to a >10,000-fold decrease in virus loads; the lungs of the treated mice were free of infiltrating leukocytes; there was no sign of pulmonary inflammation and the mice did not experience the decrease in body weight that normally occurs in untreated or control vector-treated mice. Delivery of the decoy by a lentiviral vector was as effective and appeared to be even more durable. The vectors were well-tolerated; they did not disturb myeloid or lymphoid cell populations and did not cause T cell activation or increased levels of proinflammatory cytokines in the sera. The vectors established protection against a broad range of SARS-CoV-2 variants including the recent Omicron subvariants BA.2.75, BA.4/5 and BQ.1. Protection was strongest against virus with the parental D614G spike protein and somewhat less effective against the BA.2 variant, an effect that was probably due to the relative decrease in spike protein affinity for ACE2^16^.

Administration of the decoy-expressing AAV2.retro and AAV6.2 vectors by i.n. instillation suppressed virus replication in the mice for at least 30 days. The effectiveness of i.n. administration of the AAV2.retro vector, which was somewhat greater than the AAV6.2 vector, was surprising as its capsid was selected for high efficiency transduction of the CNS and retrograde transport in neurons^34,35^; its tropism for the respiratory tract has not, to our knowledge, been previously described. The tropism of the AAV2.retro vector for lung and neuronal cells could be clinically advantageous as a means to suppress SARS-CoV-2 replication in respiratory and olfactory tissues. SARS-CoV-2 infection of ACE2 K18 Tg results in high virus loads in the brain which was suppressed by administration of the decoy-expressing AAV2.retro vector (not shown). Imaging of mice following administration of a luciferase-expressing AAV2.retro vector showed transduction of the olfactory region of the brain (not shown).

In a previous report, Sims *et al*., used an AAV-expressed high affinity ACE2 decoy to protect mice from SARS-CoV-2 infection^53^. In that study, i.n. administration of a decoy-expressing AAVhu.68 vector caused at 30-fold decrease in Wuhan-Hu-1 SARS-CoV-2 virus load 7-dpi but at 4-dpi, close to the time at which virus loads peak, had no significant effect on virus loads. In contrast, we found that i.n. or i.m. administration of decoy-expressing AAV2.retro or AAV6.2 vectors decreased virus loads 10,000-100,000-fold at the time of peak virus load. The increased effectiveness of the therapy in our study does not appear to have resulted from differences in increased neutralizing activity of the decoys which appeared to be similar in both studies (IC_50_s (37 ng/ml vs 20 ng/ml^29^) or differences in vector dosage which also appeared to be similar. A potential explanation is that of more efficient transduction of respiratory tract cells by the AAV2.retro and AAV6.2 vectors.

The effectiveness of i.m. injection with the AAV2.retro vector was encouraging because clinically this route of administration may be more practical than i.n. instillation^43^. For reasons that are unclear, i.m. injection of the AAV2.retro vector was 3-logs more effective than injection by that route with the AAV6.2 vector. It is possible that the increased effectiveness of the vector was the result of retrograde transport of the AAV2.retro capsid in myocytes or simply caused by more efficient transduction of lung cells. The efficacy of i.m. injection of the vector suggests that decoy protein synthesized by transduced myocytes at the site of injection diffuses systemically, establishing a concentration in the respiratory tract sufficient to inhibit virus replication. Similarly, in the rhesus macaque SIV model, i.m. administration of AAV vectors expressing broadly neutralizing antibodies suppressed virus loads of SIV, a virus that replicates in secondary lymphoid organs^3,39^. The long-lasting suppression of SIV replication by the vector suggests that transduced terminally differentiated myocytes can produce AAV vector-encoded proteins for a period of several years. Similarly, i.m. administration of a decoy-expressing AAV vector could provide long-lived protection in humans. The protection could be more durable than that of the extended half-life monoclonal antibodies currently in clinical use^44^.

Unexpectedly, the decoy-expressing AAVs were also effective therapeutically. I.n. instillation of the vectors as late as 24 hours post-SARS-CoV-2 infection suppressed virus replication, a time course similar to what is found in the treatment of mice with highly potent neutralizing monoclonal antibody^8^. The effectiveness of the decoy therapeutically demonstrates the rapid kinetics with which the vectors transduce cells in the lung and program biosynthesis of the encoded protein. In clinical practice, monoclonal antibody therapy is effective when given several days post-infection^45^. The similarity in the timing with which the AAV vectors and monoclonal antibodies can treat mice suggests that in humans, decoy-expressing AAV might be effective up to several days post-infection, as is the case for the use of monoclonal antibodies. In mouse and hamster models, the administration of recombinant ACE2 decoy protein has previously been shown to be highly effective therapeutically^28,46^. The proteins do not require viral transduction or biosynthesis in the lung and thus are expected to act faster; however, in clinical practice, their use will require large quantities of highly pure recombinant protein which is not the case for vectored immunoprophylaxis.

Delivery of the decoy protein with a lentiviral vector was also highly effective for the establishment of vectored immunoprophylaxis. Like the AAV vectors, the lentiviral vector was most effective administered by i.n. instillation, Unlike the AAV vectors, the lentiviral vector was also effective by i.v. injection, a route that results mainly in the transduction of splenocytes, many of which are DCs^47^. Lentiviral vectored immunoprophylaxis appeared to be more durable than AAV vector, remaining fully intact through the 60-day time course. Interestingly, the level of virus load suppression intensified at the later time points, an effect that was probably the result of increased expression levels of the decoy in the in lung as demonstrated using a luciferase-expressing reporter vector. The parameters that affected the durability of protection by the two types of vectors are unclear. The AAV vectors transduced a significantly higher proportion of lung endothelial cells while the lentiviral vector high transduced a high proportion of leukocytes, many of which were DCs. Endothelial cells are mitotically active, thus diluting the AAV genome copy number over time. DCs are terminally differentiated and could remain resident in the lung for a long time.

AAV vectors are advantageous for clinical use because they are expressed long-term without integrating into the host cell genome and are replication-defective^40,48–50^. The vectors are currently in use in a large number of clinical trials for a broad range of diseases. There are over 20 AAV serotypes, each with unique tissue tropism^51^. AAV6.2 is a variant of AAV6 that contains a single point mutation (F129L) introduced to increase lung cell tropism^36^. AAV2.retro is an AAV2 variant selected for retrograde transport in the CNS^52^. Both vectors administered i.n. protected mice from infection. AAV2.retro, which has not been reported to transduce cells of the respiratory system was, unexpectedly, somewhat more effective than AAV6.2. The majority of cells transduced by both vectors were lung the epithelial cells although AAV2.retro transduced a large proportion of neutrophils and monocytes.

The decoy-expressing lentiviral vector also strong protected mice from infection. Lentiviral vectors are currently being developed for several clinical applications including CAR-T cells and SARS-CoV-2 vaccines^54,55^. The vectors offer long-term expression and are not generally subject to pre-existing immunity^56^. The protection established by i.n. administration of the decoy-expressing vector remained intact over the 60-day time-course and was somewhat longer-lasting than that of AAV vectored protection which started to wane after 30-days. The suppression of virus replication by the lentiviral vector increased somewhat towards the end of the time-course, an effect that was associated with a small increase in expression of the decoy-expressing lentiviral vector in the lung. The increased durability of lentiviral-vectored immunoprophylaxis is presumably the result of the transduction of a long-lived cell subpopulation in the lung although the identity of this subpopulation is unclear. The lentiviral vector mainly transduced lung leukocytes, many of which were DCs and monocytes that are thought to have short 2 day half-lives^57,58^ and thus unlikely to account for the durable expression. The AAV vectors transduced a higher proportion of lung epithelial cells, a cell-type which in mice, has a much longer 17 month half-life^59^. Possible explanations for the long-lasting protection provided by the lentiviral vector are that a long-lived, subpopulation of tissue resident DC or myeloid cells had been transduced or that integration of the vector allows for persistence of its genome in dividing cell subpopulations of the lung. In most clinical applications, lentiviral vectors are used to transduce cells *ex vivo* that are later re-infused, a procedure that has been generally viewed as low-risk. The safety profile of integrating lentiviral vectors for direct injection has not been fully established^60–62^.

A potential application of vectored immunoprophylaxis for SARS-CoV-2 is as a means of providing protection to immunocompromised individuals for whom vaccination is less effective. Until recently, the most effective protection available for such individuals was the AstraZeneca Evusheld cocktail, a mixture of two monoclonal antibodies formulated for slow release by intramuscular injection^44^. However, the therapy may have become less effective as a result of immunoevasion by new Omicron subvariants. Both of the monoclonal antibodies in the cocktail have decreased neutralizing titers against the Omicrons BA.1 and BA.2^5–14^ and recent findings suggest that they may be inactive against the increasingly prevalent BQ.1 and BA.2.75 subvariants^63^. This decrease in neutralizing activity contrasts with the decoy which maintains its effectiveness against BQ.1 and BA.2.75.

AAV-based vectored immunoprophylaxis was effective therapeutically in the mouse models when delivered within a 24-hour window post-infection. The short window is at least partially a function of the rapid kinetics of virus replication and clearance in the mouse model as compared to in humans. While the decoy-expressing AAV lost efficacy at later time points in the infected mouse, the loss of effect was coincident with that found for treatment with a highly potent therapeutic monoclonal antibody. Monoclonal antibody therapy has been found to lessen disease symptoms when given to patients several days post-infection^45^. If the comparison pertains in humans, the window for which the AAV therapy remains effective in humans might be comparable to that for monoclonal antibody therapy.

Vectored immunoprophylaxis could be valuable in the case of a future pandemic spurred by the zoonosis of a novel coronavirus. Species such as bats and pangolins harbor large numbers of coronaviruses with the ability to use hACE2^64,65^. In the case of zoonosis of a coronavirus that used ACE2 as its entry receptor, the decoy-expressing vectors reported here would be ready as an off-the-shelf agent available prior to the production of a vaccine. The protection established by the vectors is more rapid than of vaccine as it does not require the induction of an immune response. In the case of zoonosis of a virus that used a receptor other than ACE2 or if a novel SARS-CoV-2 variant were to emerge that switched its receptor usage, the decoy receptor approach could also be applicable. This would involve the identification of the entry receptor for the novel virus and the construction of a soluble form of the protein to serve as a decoy. While a switch in receptor usage is possible, it has not happened to date despite strong selective pressure on the viral spike protein to alter its amino acid sequence.

## STAR Methods

### Resource Availability

#### Lead Contact

Further information and requests for resources and reagents should be directed to and will be fulfilled by the Lead Contact, Nathaniel R. Landau.

#### Materials Availability

All unique DNA constructs, proteins and pseudotyped virus generated in this study are available from the Lead Contact upon request.

#### Data and Code Availability

- The data used in this study are available upon request from the lead contact.
- This paper does not report original code.
- Any additional information required to reanalyze the data reported in this paper is available from the lead contact upon request.

### Experimental Model and Subject Details

#### Cells

293T cells were cultured in DMEM/10% FBS. Clonal cell-lines CHME3.ACE2 and A549.ACE2 were established by lipofection of CHME3 and A549 cells with plenti.ACE2^6^ using lipofectamine2000 (Invitrogen). The cells were selected in 1 μg/ml puromycin and single cell clones were evaluated by flow cytometry for high ACE2 expression. CHME3.ACE2, A549.ACE2 and ACE2.TMPRSS2.Vero E6 cells were maintained in medium with 1 μg/ml puromycin. hSABCi-NS1.1 cells were differentiated in air-liquid interface cultures in transwell dishes at 1.5 × 10^5^ cells/well. The cells were plated onto inserts coated with human type IV collagen (Sigma) in PneumaCult Ex Plus medium (Stemcell Technologies). The cells were cultured at 37 °C under 5% CO_2_. The medium in the upper and lower chambers was changed one day after plating and the medium in the lower chamber was replaced every 2 days. The medium in the upper chamber was removed the apical surface was washed with PBS weekly for 2 weeks.

#### Mice

C57BL/6 mice were from Taconic. Balb/c and hACE2 K18 Tg [B6.Cg-Tg(K18-ACE2)2Prlmn/J] were from The Jackson Laboratory. Animal use and care was approved by the NYU Langone Health Institutional Animal Care and Use Committee (#170304) according to the standards set by the Animal Welfare Act.

#### Plasmids

The expression vectors used for the production of AAV vectors were AAV.retro Rep/Cap2 (Addgene 81070), Rep/Cap6 (Addgene 110770), pAdDeltaF6 (Addgene 112867) and pAAV-CAG-tdTomato. Rep/Cap6.2 expression plasmid was generated by overlap extension PCR using Rep/Cap6 template to introduce the F129L mutation. The amplicon was cloned into the EcoR-I and Nru-I sites of Rep/Cap6. To construct GFP/nanoluciferase-expressing AAV vectors pAAV-GFP.nLuc, pAAV-CAG-ACE2.1mb.nLuc and pAAV-CAG-ACE2.1mb, DNA fragments encoding GFP.nLuc, ACE2.1mb.nLuc and ACE2.1mb were amplified by PCR and joined by overlap extension PCR using primers containing Kpn-I and EcoR-I sites. The insert was removed from pAAV-CAG-tdTomato by cleavage with Kpn-I and EcoR-I and replaced with similarly cleaved amplicon. Decoy expression vector pcACE2.1mb has been previously described^21^. Expression plasmids used to produce lentiviral pseudotypes were pMDL, pcVSV.G, pRSV.Rev, the lentiviral transfer vector plenti.GFP.nLuc^6^. Expression vectors for the SARS-CoV-2 D614G, Omicron BA.1, Omicron BA.2 spike proteins have been previously described^9,12,21^. Expression vectors for the Omicron BA.4/5, BA.2.75 and BQ.1 spike proteins were constructed by overlap extension PCR mutagenesis using the D614G^5^ spike protein plasmid as template and cloned into pcDNA6.

### Method Details

#### AAV vector stocks

AAV vector stocks were produced by cotransfection of 293T cells with pAAV-CAG-ACE2.1mb, pAdDeltaF6 and AAV.retro RepCap2 or Rep/Cap6.2 at a ratio of 25:25:30 by the calcium phosphate method. Virus-containing supernatant was harvested 2 days post-transfection. The virus was concentrated by ultracentrifugation on 40% sucrose cushion at 4°C for 16 hours at 30,000 x g, resuspended in PBS and concentrated on an Amicon Ultra Centrifugal Filter Unit (Millipore). Virus titers were measured by RT-qPCR with a primer pair and probe that hybridized to the AAV2 ITR sequences^66^.

#### SARS-CoV-2 virus stocks

SARS-CoV-2 WA1/2020 (BEI Resources, NR-52281), Omicron BA.1 (BEI Resources, NR-56461), BA.2 (BEI Resources, NR-56781) and BA.5 virus (BEI Resources, NR-58616) stocks were prepared by infection of ACE2.TMPRSS2.Vero E6 cells at an MOI=0.05 (BEI Resources, NR-56781). 2 hours post-infection, the input virus was removed and a day later, the virus-containing supernatant was filtered through a 0.45 μm filter, concentrated on an Amicon Ultra Centrifugal Filter Unit (Millipore) and frozen at −80°C in aliquots.

#### Decoy pull-down

CHME3.ACE2 or A549.ACE2 cells (1 × 10^6^) were infected with AAV2.retro, AAV6.2-ACE2.1mb or plenti.ACE2.1mb at MOI=0.5. The virus was removed the following day and the supernatant was harvested 3 days later. The decoy protein was pulled-down by a 1-hour incubation with 30 μl nickel-nitrilotriacetic acid-agarose beads (QIAGEN) and eluted in Laemmle loading buffer. The protein was then analyzed on an immunoblot probed with anti-His antibody and horseradish peroxidase (HRP)-conjugated goat anti-mouse IgG secondary antibody (Sigma-Aldrich). The signals were developed with Immobilon Crescendo Western HRP Substrate (Millipore) and visualized on an iBright imaging system (Invitrogen).

#### Pseudotype neutralization assay

D614G, BA.1, BA.2, BA.2.75, BA.4/5 and BQ.1 spike protein-pseudotyped lentiviruses were generated by co-transfection of 293T cells with pMDL, pRSV.Rev, plenti.GFP.nLuc and spike protein expression vector and normalized for reverse transcriptase activity as previously described^5^. CHME3.ACE2 and A549.ACE2 cells were transduced with serially diluted decoy-expressing AAV or lentiviral vector. The medium was removed the following day and the cells were challenged with pseudotyped viruses (MOI=0.2). Luciferase activity in duplicate samples was measured 2-dpi in an Envision 2103 microplate luminometer (PerkinElmer).

#### Flow cytometry

GFP-expressing AAV or lentiviral vectors were injected into hACE2 K18 Tg (AAV) or SAMHD1 Knockout mice (lentivirus) via i.n. injection. At 3-dpi, the lungs were homogenized in ACK buffer and the cells were disaggregated by a 30-minute treatment with 1.5 mg/mL collagenase and 0.1 mg/mL DNase followed by passage through a 0.22 μm mesh. The cells were blocked with anti-CD16/CD32 and stained with Alexa 700-anti-CD45, PerCP-Cy5.5-anti-F4/80, APC-Cy7-SiglecF, PE-Cy7-anti-CD11c, PE-Cy7-anti-CD19, APC-anti-CD3, Pacblue-anti-CD11b, PE-Cy5.5-anti-CD62L, APC-anti-CD14 and PE-Ly6C/Ly6G (Gr1) (BioLegend) and analyzed on a Beckman CytoFLEX flow cytometer using with FlowJo software. Cell-types were classified as epithelial (CD45-), alveolar macrophages (CD45+, F4/80+, SiglecF+), interstitial macrophages (CD45+, F4/80+, SiglecF-), DCs (CD45+, F4/80-, CD11c+), T cells (CD45+, CD3+), B cells (CD45+, CD19+), monocytes (CD45+, CD11b+, CD14+) and neutrophils (CD45+, CD62L+, Ly6C/Ly6G+).

#### Anti-inflammatory cytokine assay

hACE2 K18 Tg were administrated 1 × 10^12^ vg decoy-expressing AAV or 5 x 10^6^ IU decoy-expressing lentiviral vector. Mice treated with AAV or lentiviral vector were challenged 3- or 7-days later, respectively, with 2 x 10^4^ PFU SARS-CoV-2 WA1/2020. Sera were harvested 3-dpi and IFN-γ, MCP-1, TNF-α, IL-10, IL-12 and IL-6 were measured by cytokine bead array using the BD Cytometric Bead Array Mouse Inflammation Kit (BD Biosciences).

#### *In vivo* and *in vitro* luciferase assays

Balb/c or SAMHD1 knockout mice were administered AAV2.retro or AAV6.2-ACE2.1mb.nLuc (1× 10^12^ vg) or plenti.GFP.nLuc (5 x 10^6^ IU) by i.n. instillation. The mice were imaged over 30 days by the injection of 100 μl 1:40 diluted Nano-Glo substrate (Nanolight) and visualization on an IVIS Lumina III XR (PerkinElmer). To measure luciferase activity in the tissues, organs were harvested and homogenized in lysing matrix D tubes with a FastPrep-24 5G homogenizer (MP Biomedicals). Nano-Glo Luciferase Assay Reagent (Nanolight) was added and luminescence was measured on an Envision 2103 plate reader (PerkinElmer).

#### Live virus infection of cell-lines

CHME3.ACE2, A549.ACE2 and hSABCi-NS1.1 cells (2 x 10^5^) were infected with AAV2.retro or AAV6.2-ACE2.1mb at MOI=0.5. The medium was replaced 1-dpi and the following day, the cells were infected with SARS-CoV-2 at MOI=0.01. The cultures were lysed 2-dpi after which RNA was prepared and cell-associated viral RNA copies were quantified by reverse transcriptase RT-qPCR. Absolute RNA copy numbers were calculated using a standard curve generated by the analysis of a serially diluted *in vitro* transcribed synthetic subgenomic viral RNA using the 2-ΔΔCT method.

#### Prophylactic and therapeutic administration of decoy-expressing vectors

For prophylaxis experiment, 6-8 weeks old hACE2 K18 Tg or Balb/c mice were anesthetized with isoflurane or ketamine–xylazine cocktail and injected with 80 μl (i.n. or i.v. or i.m.) (1 × 10^12^ vg) of AAV2.retro or AAV6.2-decoy or 5 x 10^6^ IU of plenti.ACE2.1mb. After 1-30 days (AAV) and 1-60 days (lentivirus vector) of infection, the mice were infected i.n. with 2 x 10^4^ plaque-forming unit (PFU) of SARS-CoV-2 WA1/2020 (hACE2 K18 Tg) or Omicron BA.1 or BA.2 or BA.5 (Balb/c). At 2-dpi (Omicron) or 3-dpi (SARS-CoV-2 WA1/2020), the mice were sacrificed and RNA was prepared from 200 μl of lung lysate using the Quick-RNA MiniPrep kit (Zymo Research). For therapeutic testing, hACE2 K18 Tg were infected i.n. with 2 x 10^4^ PFU SARS-CoV-2 WA1/2020. The mice were infected 0-48 hours post-infection i.n. with 80 μl (1 × 10^12^ vg) of AAV2.retro or AAV6.2-decoy. 3-dpi (SARS-CoV-2 WA1/2020), the mice were sacrificed and RNA was prepared from 200 μl of the lung lysate using a Quick-RNA MiniPrep kit.

#### Virus loads

SARS-CoV-2 E gene subgenomic RNA levels were measured by reverse transcriptase RT-qPCR with a TaqMan probe. Lung RNA was mixed with TaqMan Fast Virus 1-step Master Mix (Applied Biosystems), 10 mM forward and reverse primers, and 2 mM probe. PCR cycles were 5 minutes at 50°C, 20 seconds at 95°C, 40 cycles of 3 seconds at 95°C, 3 seconds at 60°C). E gene subgenomic RNA copies were measured using forward primer subgenomic F (CGATCTCTTGTAGATCTGTTCTC), reverse primer E Sarbeco R and probe E Sarbeco P1)^67,68^. Tissue analyses were normalized to GAPDH mRNA copies measured using the mouse GAPDH.forward (CAATGTGTCCGTCGTGGATCT) and mouse GAPDH.reverse (GTCCTCAGTGTAGCCCAAGATG) with mouse GAPDH probe (FAM-CGTGCCGCCTGGAGAAACCTGCC-BHQ) or human GAPDH.forward (GTCTCCTCTGACTTCAACAGCG) and human GAPDH.reverse (ACCACCCTGTTGCTGTAGCCAA) with human GAPDH probe (FAM-TAGGAAGGACAGGCAAC-IBFQ). Absolute RNA copy numbers were calculated using a standard curve generated by the analysis of a serially diluted *in vitro* transcribed synthetic subgenomic viral RNA containing the E gene sequence (2019-nCoV_E Positive Control, IDT: 10006896) using the 2-ΔΔCT method.

#### Histology

The lungs of SARS-CoV-2-infected mice were harvested 3-dpi. The tissue was fixed in 10% neutral buffered formalin for 72 hours at room temperature and then processed through graded ethanol, xylene and into paraffin in a Leica Peloris automated processor. 5 μm paraffin-embedded sections were deparaffinized and stained with hematoxylin (Leica, 3801575) and eosin (Leica, 3801619) on a Leica ST5020 automated histochemical strainer. Slides were scanned at 40× magnification on a Leica AT2 whole slide scanner and the images were transferred to the NYULH Omero web-accessible image database.

## Statistical Analysis

Statistical significance was determined by Kruskal-Wallis test with post hoc Dunn’s test. Significance was calculated based on two-sided testing and is shown in the figures as the mean ± SD with confidence intervals listed as *p<0.05, **p<0.01, ***p<0.001, ****p<0.0001.

## Study Approval

Animal procedures were performed with the written approval of the NYU Animal Research Committee in accordance with all federal, state, and local guidelines.

## Acknowledgements

We thank Meike Dittman for ACE2.TMPRSS2.Vero E6 cells and the NYU Langone Laura and the Isaac Perlmutter Cancer Center Experimental Pathology Research Laboratory for histology. We thank David J. Simon (Weill Cornell Medicine) for providing AAV plasmids.

## Author Contributions

T.T. and N.R.L. designed the experiments. T.T., B.M.D. and J.M. carried out the experiments. T.T. and N.R.L. wrote the manuscript. N.R.L. supervised the study and revised the manuscript.

---

**Figure S1.**
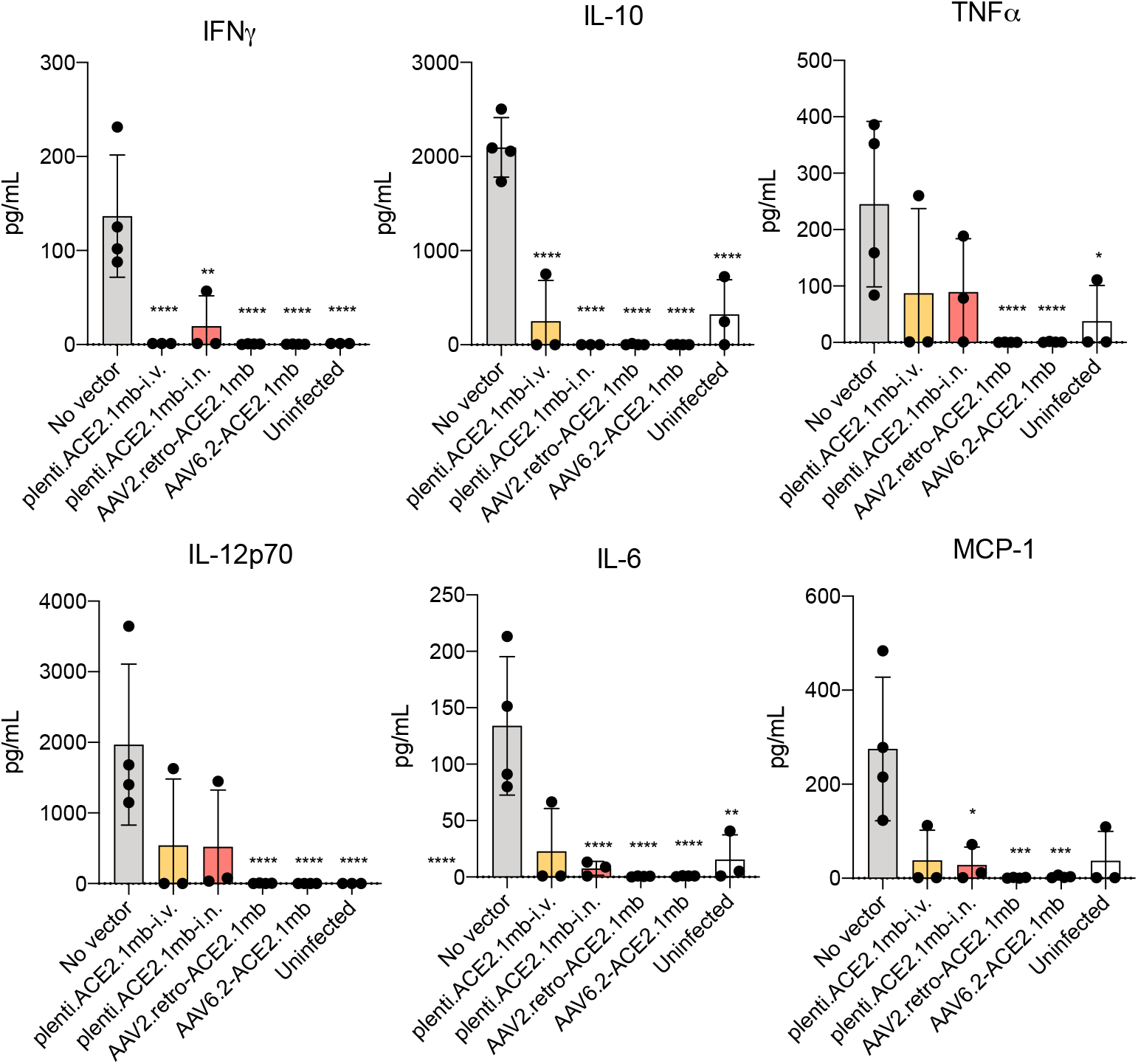
Intranasal injected AAV2.retro, AAV6.2-decoys and lentivirus-based ACE2.1mb didn’t induce any inflammatory cytokine secretion. Mice (n=4) were treated with the decoy-expressing AAV vectors (1 × 10^12^ IU) or decoy-expressing lentiviral vector (5 x 10^6^ IU). After 3 (AAV) or 7days (lentiviral vector), the mice were challenged with SARS-CoV-2 WA1/2020 and 3-dpi the levels of IFNγ, TNFα, IL-10, IL-6, MCP-1 and IL-12p70 in lung were measured by cytokine beads array. The Y-axis shows the concentration of each cytokine. The experiment was done twice with similar results. Confidence intervals are shown as the mean ± SD. *P ≤ 0.05, **P ≤ 0.01, ***P≤0.001, ****P≤0.0001.

**Figure S2.**
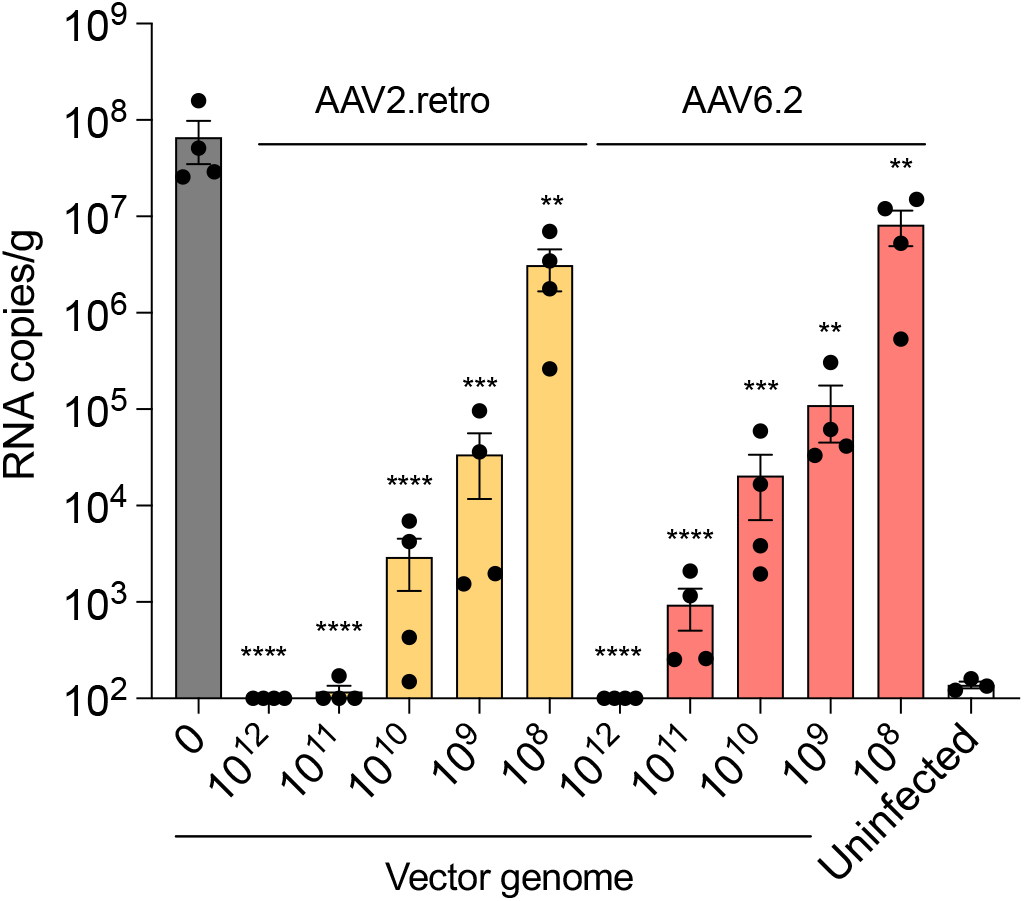
AAV-Decoy protected mice from SARS-CoV-2 infection. Different doses of decoy-expressing AAV vectors (1 × 10^12^, 1 × 10^11^, 1 × 10^10^, 1 × 10^9^, 1 × 10^8^ vg) were administered to hACE2 K18 Tg (n=4) by i.n. instillation. 3 days post-AAV injection, mice were challenged with 1 × 10^4^ PFU of SARS-CoV-2 WA1/2020. At 3-dpi, subgenomic viral E gene RNA in the lung were quantified. Confidence intervals are shown as the mean ± SD. **P ≤ 0.01, ***P≤0.001, ****P≤0.0001. The experiment was done twice with similar results.

**Figure S3.**
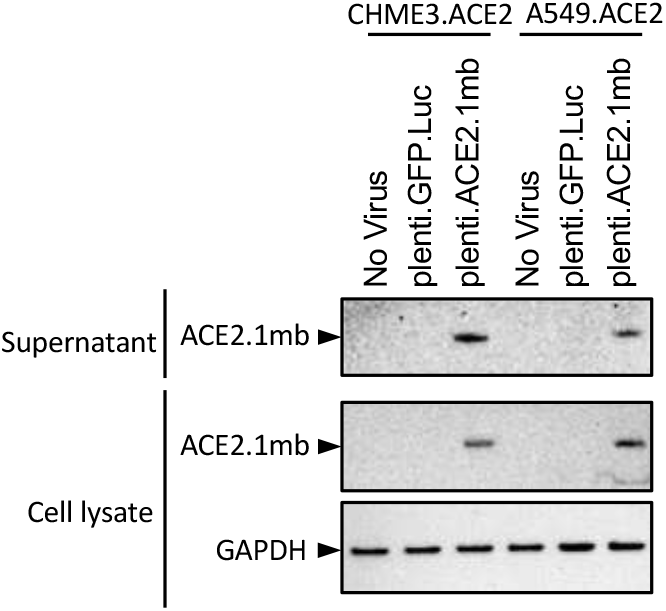
Lentivirus-based ACE2.1mb protected mice from SARS-CoV-2 infection. CHME3.ACE2 and A549.ACE2 cells were transduced with decoy-expressing lentiviral vectors at an MOI of 0.5. 3-dpi, decoy protein secreted into the supernatant was pulled-down on NTA beads and bead-bound decoy protein was detected on an immunoblot probed with His-tag antibody. Decoy protein in the cell lysates is shown below with GAPDH as a loading control.

**Figure S4.**
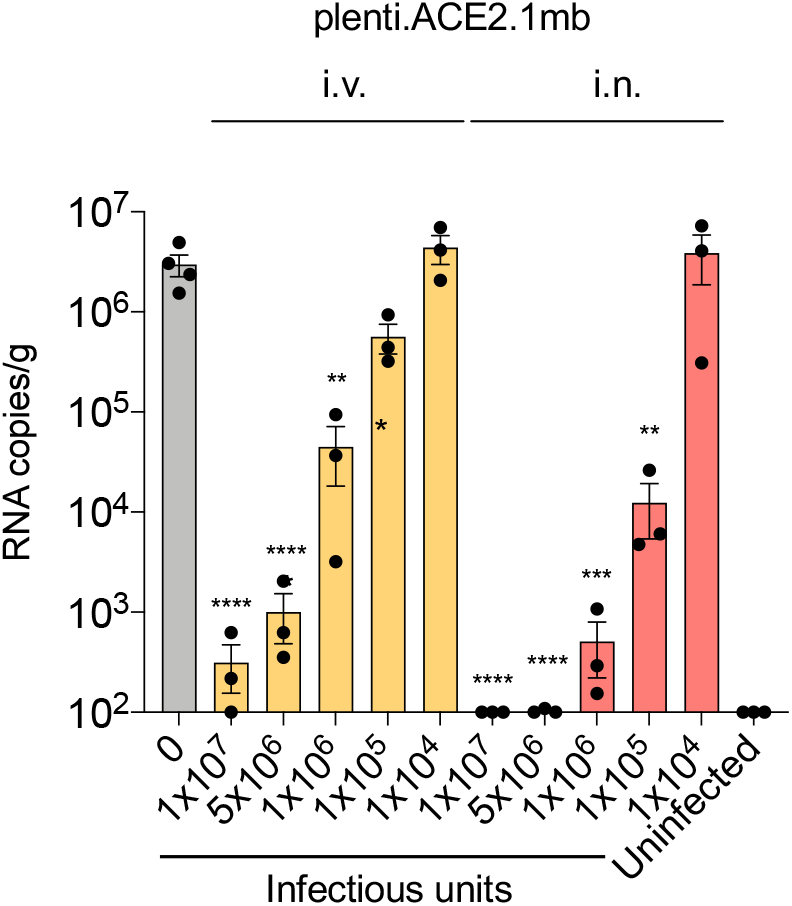
Lentivirus-based ACE2.1mb protect mice from SARS-CoV-2 infection. hACE2 K18 Tg mice (n=3) were injected i.v. or i.n. with different amounts of decoy-expressing lentiviral vectors (1 × 10^7^, 1 × 10^6^, 1 × 10^5^, 1 × 10^4^ IU). One week later, the mice were challenged with 1 × 10^4^ PFU SARS-COV-2. At 3-dpi, lung subgenomic viral E RNA was quantified by RT-PCR. Confidence intervals are shown as the mean ± SD. **P ≤ 0.01, ***P≤0.001, ****P≤0.0001.

## References

1. Johnson, P.R., Schnepp, B.C., Zhang, J., Connell, M.J., Greene, S.M., Yuste, E., Desrosiers, R.C., and Clark, K.R. (2009). Vector-mediated gene transfer engenders long-lived neutralizing activity and protection against SIV infection in monkeys. Nat Med 15, 901–906. 10.1038/nm.1967.

2. Gardner, M.R. (2020). Promise and Progress of an HIV-1 Cure by Adeno-Associated Virus Vector Delivery of Anti-HIV-1 Biologics. Front Cell Infect Microbiol 10, 176. 10.3389/fcimb.2020.00176.

3. Gardner, M.R., Kattenhorn, L.M., Kondur, H.R., von Schaewen, M., Dorfman, T., Chiang, J.J., Haworth, K.G., Decker, J.M., Alpert, M.D., Bailey, C.C., et al. (2015). AAV-expressed eCD4-Ig provides durable protection from multiple SHIV challenges. Nature 519, 87–91. 10.1038/nature14264.

4. Taylor, P.C., Adams, A.C., Hufford, M.M., de la Torre, I., Winthrop, K., and Gottlieb, R.L. (2021). Neutralizing monoclonal antibodies for treatment of COVID-19. Nat Rev Immunol 21, 382–393. 10.1038/s41577-021-00542-x.

5. VanBlargan, L.A., Errico, J.M., Halfmann, P.J., Zost, S.J., Crowe, J.E., Jr., Purcell, L.A., Kawaoka, Y., Corti, D., Fremont, D.H., and Diamond, M.S. (2022). An infectious SARS-CoV-2 B.1.1.529 Omicron virus escapes neutralization by therapeutic monoclonal antibodies. Nat Med. 10.1038/s41591-021-01678-y.

6. Liu, L., Iketani, S., Guo, Y., Chan, J.F.W., Wang, M., Liu, L., Luo, Y., Chu, H., Huang, Y., Nair, M.S., et al. (2022). Striking antibody evasion manifested by the Omicron variant of SARS-CoV-2. Nature 602, 676–681. 10.1038/s41586-021-04388-0.

7. Planas, D., Saunders, N., Maes, P., Guivel-Benhassine, F., Planchais, C., Buchrieser, J., Bolland, W.-H., Porrot, F., Staropoli, I., Lemoine, F., et al. (2021). Considerable escape of SARS-CoV-2 Omicron to antibody neutralization. Nature. 10.1038/s41586-021-04389-z.

8. Dragoni, F., Schiaroli, E., Micheli, V., Fiaschi, L., Lai, A., Zehender, G., Rossetti, B., Gismondo, M.R., Francisci, D., Zazzi, M., and Vicenti, I. (2022). Impact of SARS-CoV-2 omicron BA.1 and delta AY.4.2 variants on the neutralization by sera of patients treated with different authorized monoclonal antibodies. Clin Microbiol Infect. 10.1016/j.cmi.2022.03.005.

9. Zhou, H., Dcosta, B.M., Landau, N.R., and Tada, T. (2022). Resistance of SARS-CoV-2 Omicron BA.1 and BA.2 Variants to Vaccine-Elicited Sera and Therapeutic Monoclonal Antibodies. Viruses 14. 10.3390/v14061334.

10. Iketani, S., Liu, L., Guo, Y., Liu, L., Chan, J.F.W., Huang, Y., Wang, M., Luo, Y., Yu, J., Chu, H., et al. (2022). Antibody evasion properties of SARS-CoV-2 Omicron sublineages. Nature 604, 553–556. 10.1038/s41586-022-04594-4.

11. Chen, R.E., Winkler, E.S., Case, J.B., Aziati, I.D., Bricker, T.L., Joshi, A., Darling, T.L., Ying, B., Errico, J.M., Shrihari, S., et al. (2021). In vivo monoclonal antibody efficacy against SARS-CoV-2 variant strains. Nature 596, 103–108. 10.1038/s41586-021-03720-y.

12. Tada, T., Zhou, H., Dcosta, B.M., Samanovic, M.I., Chivukula, V., Herati, R.S., Hubbard, S.R., Mulligan, M.J., and Landau, N.R. (2022). Increased resistance of SARS-CoV-2 Omicron variant to neutralization by vaccine-elicited and therapeutic antibodies. EBioMedicine 78, 103944. 10.1016/j.ebiom.2022.103944.

13. Diamond, M., Halfmann, P., Maemura, T., Iwatsuki-Horimoto, K., Iida, S., Kiso, M., Scheaffer, S., Darling, T., Joshi, A., Loeber, S., et al. (2021). The SARS-CoV-2 B.1.1.529 Omicron virus causes attenuated infection and disease in mice and hamsters. Res Sq. 10.21203/rs.3.rs-1211792/v1.

14. Bruel, T., Hadjadj, J., Maes, P., Planas, D., Seve, A., Staropoli, I., Guivel-Benhassine, F., Porrot, F., Bolland, W.H., Nguyen, Y., et al. (2022). Serum neutralization of SARS-CoV-2 Omicron sublineages BA.1 and BA.2 in patients receiving monoclonal antibodies. Nat Med. 10.1038/s41591-022-01792-5.

15. Touret, F., Baronti, C., Pastorino, B., Villarroel, P.M.S., Ninove, L., Nougairede, A., and de Lamballerie, X. (2022). In vitro activity of therapeutic antibodies against SARS-CoV-2 Omicron BA.1, BA.2 and BA.5. Sci Rep 12, 12609. 10.1038/s41598-022-16964-z.

16. Cao, Y., Yisimayi, A., Jian, F., Song, W., Xiao, T., Wang, L., Du, S., Wang, J., Li, Q., Chen, X., et al. (2022). BA.2.12.1, BA.4 and BA.5 escape antibodies elicited by Omicron infection. Nature 608, 593–602. 10.1038/s41586-022-04980-y.

17. Imai, M., Ito, M., Kiso, M., Yamayoshi, S., Uraki, R., Fukushi, S., Watanabe, S., Suzuki, T., Maeda, K., Sakai-Tagawa, Y., et al. (2022). Efficacy of Antiviral Agents against Omicron Subvariants BQ.1.1 and XBB. N Engl J Med. 10.1056/NEJMc2214302.

18. Czajkowsky, D.M., Hu, J., Shao, Z., and Pleass, R.J. (2012). Fc-fusion proteins: new developments and future perspectives. EMBO Mol Med 4, 1015–1028. 10.1002/emmm.201201379.

19. Hodges, T.L., Kahn, J.O., Kaplan, L.D., Groopman, J.E., Volberding, P.A., Amman, A.J., Arri, C.J., Bouvier, L.M., Mordenti, J., and Izu, A.E. (1991). Phase 1 study of recombinant human CD4-immunoglobulin G therapy of patients with AIDS and AIDS-related complex. Antimicrobial Agents and Chemotherapy 35, 2580–2586. doi:10.1128/AAC.35.12.2580.

20. Gershon, D. (1996). Genentech sheds gp120 vaccine. Nature Medicine 2, 370–371. 10.1038/nm0496-370.

21. Tada, T., Fan, C., Chen, J.S., Kaur, R., Stapleford, K.A., Gristick, H., Dcosta, B.M., Wilen, C.B., Nimigean, C.M., and Landau, N.R. (2020). An ACE2 Microbody Containing a Single Immunoglobulin Fc Domain Is a Potent Inhibitor of SARS-CoV-2. Cell Rep 33, 108528. 10.1016/j.celrep.2020.108528.

22. Chan, K.K., Dorosky, D., Sharma, P., Abbasi, S.A., Dye, J.M., Kranz, D.M., Herbert, A.S., and Procko, E. (2020). Engineering human ACE2 to optimize binding to the spike protein of SARS coronavirus 2. Science 369, 1261–1265. 10.1126/science.abc0870.

23. Jing, W., and Procko, E. (2021). ACE2-based decoy receptors for SARS coronavirus 2. Proteins 39, 1065–1078. 10.1002/prot.26140.

24. Zhang, L., Narayanan, K.K., Cooper, L., Chan, K.K., Skeeters, S.S., Devlin, C.A., Aguhob, A., Shirley, K., Rong, L., Rehman, J., et al. (2022). An ACE2 decoy can be administered by inhalation and potently targets omicron variants of SARS-CoV-2. EMBO Molecular Medicine 14, e16109. https://doi.org/10.15252/emmm.202216109.

25. Linsky, T.W., Vergara, R., Codina, N., Nelson, J.W., Walker, M.J., Su, W., Barnes, C.O., Hsiang, T.Y., Esser-Nobis, K., Yu, K., et al. (2020). De novo design of potent and resilient hACE2 decoys to neutralize SARS-CoV-2. Science 370, 1208–1214. 10.1126/science.abe0075.

26. Higuchi, Y., Suzuki, T., Arimori, T., Ikemura, N., Mihara, E., Kirita, Y., Ohgitani, E., Mazda, O., Motooka, D., Nakamura, S., et al. (2021). Engineered ACE2 receptor therapy overcomes mutational escape of SARS-CoV-2. Nature Communications 12, 3802. 10.1038/s41467-021-24013-y.

27. Case, J.B., Rothlauf, P.W., Chen, R.E., Liu, Z., Zhao, H., Kim, A.S., Bloyet, L.M., Zeng, Q., Tahan, S., Droit, L., et al. (2020). Neutralizing Antibody and Soluble ACE2 Inhibition of a Replication-Competent VSV-SARS-CoV-2 and a Clinical Isolate of SARS-CoV-2. Cell Host Microbe. 10.1016/j.chom.2020.06.021.

28. Zhang, L., Dutta, S., Xiong, S., Chan, M., Chan, K.K., Fan, T.M., Bailey, K.L., Lindeblad, M., Cooper, L.M., Rong, L., et al. (2022). Engineered ACE2 decoy mitigates lung injury and death induced by SARS-CoV-2 variants. Nat Chem Biol 18, 342–351. 10.1038/s41589-021-00965-6.

29. Tada, T., Dcosta, B.M., Zhou, H., and Landau, N.R. (2023). ACE2 Receptor Decoy is a Potent Prophylactic and Therapeutic for SARS-CoV-2. bioRxiv, 2022.2012.2031.522401. 10.1101/2022.12.31.522401.

30. Guy, J.L., Jackson, R.M., Jensen, H.A., Hooper, N.M., and Turner, A.J. (2005). Identification of critical active-site residues in angiotensin-converting enzyme-2 (ACE2) by site-directed mutagenesis. FEBS J 272, 3512–3520. 10.1111/j.1742-4658.2005.04756.x.

31. Meyer-Berg, H., Zhou Yang, L., Pilar de Lucas, M., Zambrano, A., Hyde, S.C., and Gill, D.R. (2020). Identification of AAV serotypes for lung gene therapy in human embryonic stem cell-derived lung organoids. Stem Cell Res Ther 11, 448. 10.1186/s13287-020-01950-x.

32. van Lieshout, L.P., Domm, J.M., and Wootton, S.K. (2019). AAV-Mediated Gene Delivery to the Lung. Methods Mol Biol 1950, 361–372. 10.1007/978-1-4939-9139-6_21.

33. Halbert, C.L., Allen, J.M., and Miller, A.D. (2001). Adeno-associated virus type 6 (AAV6) vectors mediate efficient transduction of airway epithelial cells in mouse lungs compared to that of AAV2 vectors. J Virol 75, 6615–6624. 10.1128/JVI.75.14.6615-6624.2001.

34. Tervo, D.G., Hwang, B.Y., Viswanathan, S., Gaj, T., Lavzin, M., Ritola, K.D., Lindo, S., Michael, S., Kuleshova, E., Ojala, D., et al. (2016). A Designer AAV Variant Permits Efficient Retrograde Access to Projection Neurons. Neuron 92, 372–382. 10.1016/j.neuron.2016.09.021.

35. Zhang, Y., Wang, J., Li, J., Chen, Y., Sun, J., Lu, Z., Li, Y., and Liu, T. (2022). Functional analysis of mutations endowing rAAV2-retro with retrograde tracing capacity. Neurosci Lett 784, 136746. 10.1016/j.neulet.2022.136746.

36. Limberis, M.P., Vandenberghe, L.H., Zhang, L., Pickles, R.J., and Wilson, J.M. (2009). Transduction efficiencies of novel AAV vectors in mouse airway epithelium in vivo and human ciliated airway epithelium in vitro. Mol Ther 17, 294–301. 10.1038/mt.2008.261.

37. Halfmann, P.J., Iida, S., Iwatsuki-Horimoto, K., Maemura, T., Kiso, M., Scheaffer, S.M., Darling, T.L., Joshi, A., Loeber, S., Singh, G., et al. (2022). SARS-CoV-2 Omicron virus causes attenuated disease in mice and hamsters. Nature 603, 687–692. 10.1038/s41586-022-04441-6.

38. Zhang, Y.N., Zhang, Z.R., Zhang, H.Q., Li, N., Zhang, Q.Y., Li, X.D., Deng, C.L., Deng, F., Shen, S., Zhu, B., and Zhang, B. (2022). Different pathogenesis of SARS-CoV-2 Omicron variant in wild-type laboratory mice and hamsters. Signal Transduct Target Ther 7, 62. 10.1038/s41392-022-00930-2.

39. Spitsin, S., Schnepp, B.C., Connell, M.J., Liu, T., Dang, C.M., Pappa, V., Tustin, R., Kinder, A., Johnson, P.R., and Douglas, S.D. (2020). Protection against SIV in Rhesus Macaques Using Albumin and CD4-Based Vector-Mediated Gene Transfer. Molecular therapy. Methods & clinical development 17, 1088–1096. 10.1016/j.omtm.2020.04.019.

40. Wang, D., Tai, P.W.L., and Gao, G. (2019). Adeno-associated virus vector as a platform for gene therapy delivery. Nature Reviews Drug Discovery 18, 358–378. 10.1038/s41573-019-0012-9.

41. Mingozzi, F., and High, K.A. (2013). Immune responses to AAV vectors: overcoming barriers to successful gene therapy. Blood 122, 23–36. 10.1182/blood-2013-01-306647.

42. Desch, A.N., Henson, P.M., and Jakubzick, C.V. (2013). Pulmonary dendritic cell development and antigen acquisition. Immunol Res 55, 178–186. 10.1007/s12026-012-8359-6.

43. Tosolini, A.P., and Sleigh, J.N. (2020). Intramuscular Delivery of Gene Therapy for Targeting the Nervous System. Front Mol Neurosci 13, 129. 10.3389/fnmol.2020.00129.

44. Levin, M.J., Ustianowski, A., De Wit, S., Launay, O., Avila, M., Templeton, A., Yuan, Y., Seegobin, S., Ellery, A., Levinson, D.J., et al. (2022). Intramuscular AZD7442 (Tixagevimab-Cilgavimab) for Prevention of Covid-19. N Engl J Med. 10.1056/NEJMoa2116620.

45. Miguez-Rey, E., Choi, D., Kim, S., Yoon, S., and Sandulescu, O. (2022). Monoclonal antibody therapies in the management of SARS-CoV-2 infection. Expert Opin Investig Drugs 31, 41–58. 10.1080/13543784.2022.2030310.

46. Sims, J.J., Lian, S., Meggersee, R.L., Kasimsetty, A., and Wilson, J.M. (2022). High activity of an affinity-matured ACE2 decoy against Omicron SARS-CoV-2 and pre-emergent coronaviruses. PLoS One 17, e0271359. 10.1371/journal.pone.0271359.

47. Tada, T., Norton, T.D., Leibowitz, R., and Landau, N.R. (2022). Directly injected lentiviral vector-based T cell vaccine protects mice against acute and chronic viral infection. JCI Insight. 10.1172/jci.insight.161598.

48. Li, C., and Samulski, R.J. (2020). Engineering adeno-associated virus vectors for gene therapy. Nat Rev Genet 21, 255–272. 10.1038/s41576-019-0205-4.

49. Martinez-Navio, J.M., Fuchs, S.P., Mendes, D.E., Rakasz, E.G., Gao, G., Lifson, J.D., and Desrosiers, R.C. (2020). Long-Term Delivery of an Anti-SIV Monoclonal Antibody With AAV. Front Immunol 11, 449. 10.3389/fimmu.2020.00449.

50. Naso, M.F., Tomkowicz, B., Perry, W.L., 3rd, and Strohl, W.R. (2017). Adeno-Associated Virus (AAV) as a Vector for Gene Therapy. BioDrugs 31, 317–334. 10.1007/s40259-017-0234-5.

51. Wu, Z., Asokan, A., and Samulski, R.J. (2006). Adeno-associated virus serotypes: vector toolkit for human gene therapy. Mol Ther 14, 316–327. 10.1016/j.ymthe.2006.05.009.

52. Weiss, A.R., Liguore, W.A., Domire, J.S., Button, D., and McBride, J.L. (2020). Intra-striatal AAV2.retro administration leads to extensive retrograde transport in the rhesus macaque brain: implications for disease modeling and therapeutic development. Sci Rep 10, 6970. 10.1038/s41598-020-63559-7.

53. Sims, J.J., Greig, J.A., Michalson, K.T., Lian, S., Martino, R.A., Meggersee, R., Turner, K.B., Nambiar, K., Dyer, C., Hinderer, C., et al. (2021). Intranasal gene therapy to prevent infection by SARS-CoV-2 variants. PLoS Pathog 17, e1009544. 10.1371/journal.ppat.1009544.

54. Lana, M.G., and Strauss, B.E. (2020). Production of Lentivirus for the Establishment of CAR-T Cells. Methods Mol Biol 2086, 61–67. 10.1007/978-1-0716-0146-4_4.

55. Lundstrom, K. (2021). Viral Vectors for COVID-19 Vaccine Development. Viruses 13. 10.3390/v13020317.

56. Ku, M.W., Authie, P., Nevo, F., Souque, P., Bourgine, M., Romano, M., Charneau, P., and Majlessi, L. (2021). Lentiviral vector induces high-quality memory T cells via dendritic cells transduction. Commun Biol 4, 713. 10.1038/s42003-021-02251-6.

57. Kamath, A.T., Henri, S., Battye, F., Tough, D.F., and Shortman, K. (2002). Developmental kinetics and lifespan of dendritic cells in mouse lymphoid organs. Blood 100, 1734–1741.

58. van Furth, R. (1989). Origin and turnover of monocytes and macrophages. Curr Top Pathol 79, 125–150.

59. Rawlins, E.L., and Hogan, B.L. (2008). Ciliated epithelial cell lifespan in the mouse trachea and lung. Am J Physiol Lung Cell Mol Physiol 295, L231–234. 10.1152/ajplung.90209.2008.

60. Mehrabadi, A.Z., Ranjbar, R., Farzanehpour, M., Shahriary, A., Dorostkar, R., Hamidinejad, M.A., and Ghaleh, H.E.G. (2022). Therapeutic potential of CAR T cell in malignancies: A scoping review. Biomed Pharmacother 146, 112512. 10.1016/j.biopha.2021.112512.

61. Mohanty, R., Chowdhury, C.R., Arega, S., Sen, P., Ganguly, P., and Ganguly, N. (2019). CAR T cell therapy: A new era for cancer treatment (Review). Oncol Rep 42, 2183–2195. 10.3892/or.2019.7335.

62. Sterner, R.C., and Sterner, R.M. (2021). CAR-T cell therapy: current limitations and potential strategies. Blood Cancer Journal 11, 69. 10.1038/s41408-021-00459-7.

63. Akerman, A., Milogiannakis, V., Jean, T., Esneu, C., Silva, M.R., Ison, T., Fitcher, C., Lopez, J.A., Chandra, D., Naing, Z., et al. (2022). Emergence and antibody evasion of BQ and BA.2.75 SARS-CoV-2 sublineages in the face of maturing antibody breadth at the population level. medRxiv, 2022.2012.2006.22283000. 10.1101/2022.12.06.22283000.

64. Demogines, A., Farzan, M., and Sawyer, S.L. (2012). Evidence for ACE2-Utilizing Coronaviruses (CoVs) Related to Severe Acute Respiratory Syndrome CoV in Bats. Journal of Virology 86, 6350–6353. 10.1128/jvi.00311-12.

65. Wacharapluesadee, S., Tan, C.W., Maneeorn, P., Duengkae, P., Zhu, F., Joyjinda, Y., Kaewpom, T., Chia, W.N., Ampoot, W., Lim, B.L., et al. (2021). Evidence for SARS-CoV-2 related coronaviruses circulating in bats and pangolins in Southeast Asia. Nature Communications 12, 972. 10.1038/s41467-021-21240-1.

66. Aurnhammer, C., Haase, M., Muether, N., Hausl, M., Rauschhuber, C., Huber, I., Nitschko, H., Busch, U., Sing, A., Ehrhardt, A., and Baiker, A. (2012). Universal real-time PCR for the detection and quantification of adeno-associated virus serotype 2-derived inverted terminal repeat sequences. Hum Gene Ther Methods 23, 18–28. 10.1089/hgtb.2011.034.

67. Winkler, E.S., Bailey, A.L., Kafai, N.M., Nair, S., McCune, B.T., Yu, J., Fox, J.M., Chen, R.E., Earnest, J.T., Keeler, S.P., et al. (2020). SARS-CoV-2 infection of human ACE2-transgenic mice causes severe lung inflammation and impaired function. Nature Immunology 21, 1327–1335. 10.1038/s41590-020-0778-2.

68. Corman, V.M., Landt, O., Kaiser, M., Molenkamp, R., Meijer, A., Chu, D.K., Bleicker, T., Brunink, S., Schneider, J., Schmidt, M.L., et al. (2020). Detection of 2019 novel coronavirus (2019-nCoV) by real-time RT-PCR. Euro Surveill 25. 10.2807/1560-7917.ES.2020.25.3.2000045.

